# Deployment of endocytic machinery to periactive zones of nerve terminals is independent of active zone assembly and evoked release

**DOI:** 10.1101/2025.04.23.650151

**Authors:** Javier Emperador-Melero, Steven J. Del Signore, Kevin M. De León González, Pascal S. Kaeser, Avital A. Rodal

## Abstract

In presynaptic nerve terminals, the endocytic apparatus rapidly restores synaptic vesicles after neurotransmitter release. Many endocytic proteins localize to the periactive zone, a loosely defined area adjacent to active zones. A prevailing model posits that recruitment of these endocytic proteins to the periactive zone is activity-dependent. We here show that periactive zone targeting of endocytic proteins is largely independent of active zone machinery and synaptic activity. At mouse hippocampal synapses and *Drosophila* neuromuscular junctions, pharmacological or genetic silencing resulted in unchanged or increased levels of endocytic proteins including Dynamin, Amphiphysin, Nervous Wreck, Endophilin A, Dap160/Intersectin, PIPK1γ and AP-180. Similarly, disruption of active zone assembly via genetic ablation of active zone scaffolds at each synapse did not impair the localization of endocytic proteins. Overall, our work indicates that endocytic proteins are constitutively deployed to the periactive zone and supports the existence of independent assembly pathways for active zones and periactive zones.

## Introduction

Release of neurotransmitters relies on the coordinated function of two complementary protein machineries, each comprising many proteins. First, the active zone defines vesicular release sites and clusters voltage-gated Ca^2+^ channels of the Ca_V_2 family for rapid fusion of synaptic vesicles in response to action potentials (Emperador-Melero and Kaeser, 2020; Sudhof, 2012). Second, the endocytic apparatus restores vesicles after exocytosis for subsequent rounds of refilling and release. The endocytic machinery is enriched at the periactive zone, a region typically localized adjacent to the active zone (Haucke et al., 2011; Saheki and De Camilli, 2012; Watanabe and Boucrot, 2017). Despite the functional relationships between exocytosis and endocytosis and the spatial coupling of their respective protein machineries, the mechanisms that determine the recruitment of the endocytic apparatus to the periactive zone remain uncertain.

One model posits that deployment of endocytic machinery to the periactive zone occurs in response to synaptic activity. Multiple lines of evidence support this model. First, depolarization causes some endocytic proteins (including Dap160/Intersectin, Endophilin and Dynamin) to partially relocalize from the vesicle pool to the cytosol or plasma membrane (Bai et al., 2010; Jiang et al., 2024; Koh et al., 2007; Winther et al., 2015, 2013). Second, membrane uptake increases rapidly in response to exocytosis triggered by synaptic activity (Haucke et al., 2011; Saheki and De Camilli, 2012; Watanabe and Boucrot, 2017), indicating that the endocytic protein machinery is responsive to activity and might be recruited to the membrane in consequence. Third, activity-dependent mechanisms at synapses might parallel protein deployment for Clathrin-mediated endocytosis in non-neuronal cells, where cytosolic endocytic machinery is transiently and acutely recruited to discrete sites (Cocucci et al., 2014; Kaksonen and Roux, 2018; Taylor et al., 2011). Fourth, the distributions of Clathrin, the Clathrin adaptor γ-Adaptin and one Dynamin isoform become diffuse after blocking action potentials in neurons with impaired endocytosis (Ferguson et al., 2007; Milosevic et al., 2011; Raimondi et al., 2011).

An alternative model in which the endocytic apparatus is constitutively recruited to the periactive zone is also supported by previous work. For example, unstimulated synapses contain endocytic machinery at the periactive zone (Bloom et al., 2003; Del Signore et al., 2023; Estes et al., 1996; Gerth et al., 2017; Imoto et al., 2022; Roos and Kelly, 1999; Wahl et al., 2013; Winther et al., 2015, 2013), raising the possibility that periactive zone assembly relies on active zone proteins or other activity-independent mechanisms. Furthermore, ultrafast endocytosis requires pre-deployment of Dynamin and pre-assembly of a ring of F-actin surrounding the active zone (Watanabe et al., 2013; Imoto et al., 2022; Ogunmowo et al., 2023). In addition, endocytic machinery is required to prevent rapid synaptic depression, which occurs at a timescale that might exceed the speed of activity-dependent protein recruitment (Hua et al., 2011; Jäpel et al., 2020; Kawasaki et al., 2000). Moreover, the endocytic machinery is involved in functions that span beyond endocytosis and are more likely to depend on predeployment of machinery to the plasma membrane. These include the regulation of the exocytic fusion pore, the fusion of dense core vesicles, and the trafficking of adhesion molecules, growth factors and extracellular vesicle cargoes (Anantharam et al., 2011; Bailey et al., 1992; Blanchette et al., 2022; Fu and Huang, 2010; Moro et al., 2021; Rodal et al., 2008; Samasilp et al., 2012). Overall, it has remained uncertain whether endocytic machinery is predeployed or recruited on an as-needed basis by activity.

Here, we combine genetic and pharmacological manipulations at two model synapses to test whether the recruitment of endocytic machinery to the periactive zone depends on synaptic activity or active zone scaffolds. Our work establishes that endocytic machinery is efficiently targeted to synapses and deployed to the periactive zone when evoked synaptic activity is chronically inhibited either pharmacologically or genetically. Furthermore, multiple active zone organizers, including RIM, ELKS and its homologue Brp, Liprin-α, and Rab3 were not required for recruitment and positioning of endocytic proteins to the periactive zone. Our findings support that endocytic machinery is constitutively deployed to presynaptic nerve terminals and localizes to periactive zones independent of active zone assembly and evoked synaptic vesicle release.

## Results

### Localization of endocytic proteins after silencing or depolarization of mouse hippocampal neurons

We generated primary neurons from mouse hippocampi and modulated their activity either by silencing them chronically or by triggering exocytosis acutely (Fig. 1A). We then assessed the distribution of endocytic proteins in these conditions compared to untreated neurons. To chronically inhibit activity, we simultaneously blocked action potentials with the Na^+^ channel blocker tetrodotoxin (TTX) and Ca^2+^ entry with blockers of Ca_V_2.1 (ω-agatoxin IVA; ω-Aga) and Ca_V_2.2 (ω-conotoxin GVIA; ω-Cono) channels. We added this drug cocktail at 3 days *in vitro* (DIV) and resupplied it every three days, which results in continuous inhibition, as established before (Held et al., 2020). To acutely trigger exocytosis in neurons that were not treated with blockers, we depolarized neurons with superfusion of 50 mM KCl for 30 seconds. These two paradigms were used to test whether synaptic transmission contributes to the assembly of the periactive zone.

**Figure 1.**
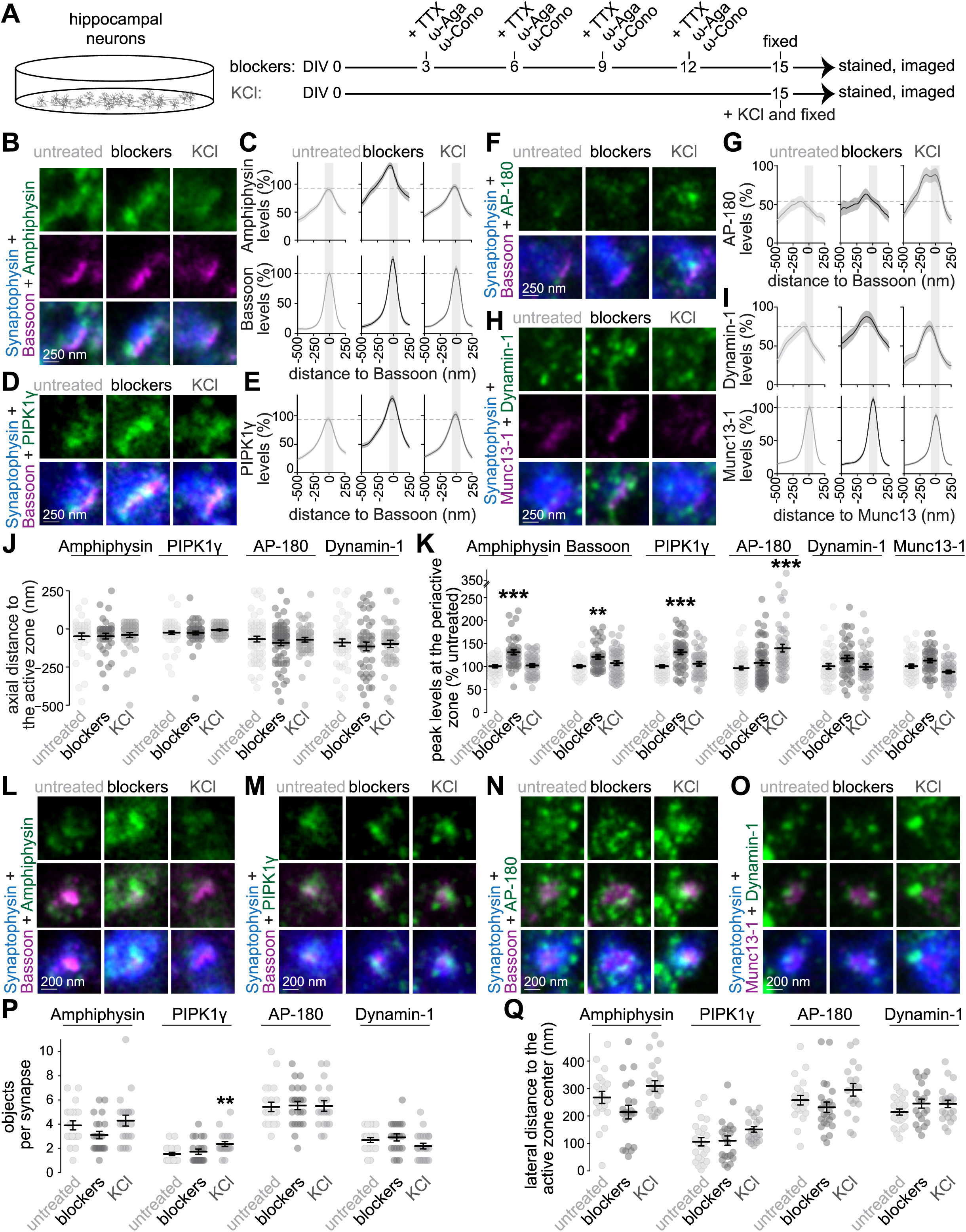
Deployment of endocytic proteins after chronic silencing or acute depolarization of mouse hippocampal neurons. (**A**) Schematic of the experiment in mouse hippocampal neurons. For chronic silencing, a cocktail (“blockers”) with tetrodotoxin (TTX, 1 μM final concentration), ω-agatoxin IVA (ω-Aga, 250 nM) and ω-conotoxin GVIA (ω-Cono, 200 nM) was added every three days starting DIV3. For acute depolarization (“KCl”), neurons were stimulated with 50 mM KCl for 30 seconds before fixation. (**B-I**) Example side-view synapses and average line profiles for neurons stained for Bassoon, Synaptophysin, and Amphiphysin (B+C), PIPK1γ (D+E), or AP-180 (F+G), or Synaptophysin, Munc13-1, and Dynamin-1 (H+I). Neurons were stained for a protein of interest (Amphiphysin, PIPK1γ, AP-180 or Dynamin-1; imaged in STED), an active zone marker (Bassoon or Munc13-1; imaged in STED), and Synaptophysin (imaged in confocal). An area of interest was positioned perpendicular to the center of the active zone marker, and synapses were aligned via the peak fluorescence of the active zone marker in the average profiles (C, E, G, I). Line profile plots were normalized to the average signal in the untreated condition. Dashed lines in C, E, G and I mark average levels in the untreated condition and grey shaded areas represent the active zone and periactive zone area; n in B, C (synapses/cultures): untreated 38/3, blockers 40/3, KCl 45/3; D, E: untreated 51/3, blockers 49/3, KCl 42/3; F, G: untreated 58/3, blockers 61/3, KCl 50/3; H, I: untreated 45/3, blockers 43/3, KCl 42/3. (**J, K**) Quantification and statistical analyses of the experiment shown in B-I, including peak-to-peak distance of the active zone marker and the protein of interest (J), and of the peak levels in the periactive zone area (K). The periactive zone area is defined as an area within 68 nm on each side of the peak of the active zone marker (grey shaded areas in C, E, G and I); n as in B-I. (**L-Q**) Example en-face synapses (L-O) and quantification of the number of objects (P) and distance of these objects to the center of the active zone marker (Q) of the experiment shown in A-K; n in P, Q (synapses/cultures): Amphiphysin, untreated 20/3, blockers 21/3, KCl 21/3; PIPK1γ, untreated 23/3, blockers 21/3, KCl 20/3; AP-180, untreated 21/3, blockers 24/3, KCl 18/3; Dynamin-1, untreated 22/3, blockers 20/3, KCl 22/3. Data are shown as mean ± SEM; *p < 0.05, **p < 0.05, ***p < 0.001 shown compared to the untreated condition determined by Kruskal-Wallis followed by Holm post hoc tests (J for Amphiphysin, PIPK1γ and AP-180; K, P, and Q), or by one-way ANOVA followed by a Tukey-Kramer post hoc test (J for Dynamin-1). For quantification of confocal images, see Fig. 1 - figure supplement 1; for a workflow of STED analyses, see Fig. 1 - figure supplement 2; for assessment of AP-180 using an independent antibody, see Fig. 1 - figure supplement 3, for additional analyses of en-face synapses, see Fig. 1 - figure supplement 4.

We fixed the neurons that were either treated with blockers (for at least 12 days) or depolarized acutely and used antibody staining to label components of the synaptic endocytic machinery. This machinery contains multiple proteins that belong to several categories including (1) BAR-domain containing proteins that sense and generate membrane invaginations, (2) phosphoinositide enzymes that regulate lipid metabolism, (3) components of Clathrin coats that sort recycled proteins, and (4) Dynamin proteins that operate in membrane fission. We labeled the BAR-domain containing protein Amphiphysin, the phosphoinositide kinase PIPK1γ, the Clathrin adaptor AP-180, and Dynamin-1 (Di Paolo et al., 2002; Koo et al., 2015; Raimondi et al., 2011; Wenk et al., 2001). We co-stained for the synaptic vesicle marker Synaptophysin, and an active zone marker (either the scaffold Bassoon or the vesicle priming protein Munc13-1, as we did before (Wong et al., 2018)). We detected signals generated by these antibodies using two complementary approaches as we described before (Emperador-Melero et al., 2024, 2021b): confocal microscopy was used to quantify protein levels in presynaptic boutons, and stimulated emission depletion (STED) microscopy was employed to assess subsynaptic protein distributions and levels at the presynaptic plasma membrane.

To estimate synaptic levels in confocal images, we quantified the average signal intensities of the proteins within regions of interest (ROIs) defined by Synaptophysin staining using a thresholding and segmentation method established before (Emperador-Melero et al., 2024, 2021b; Liu et al., 2018). The average Amphiphysin, PIPK1γ and AP-180 signal intensities were increased by up to 40% after chronic inhibition of activity, and those of Dynamin-1 were unaffected (Fig. 1 - figure supplement 1A+B). Enhanced levels in this condition were also present for the vesicle protein Synaptophysin and for Munc13-1 and may reflect a homeostatic increase of presynaptic material in response to chronic silencing. Hence, blocking action potentials and Ca^2+^ entry did not decrease the targeting of endocytic proteins to synapses. Acute depolarization enhanced the signal intensity of AP-180 by ∼30%, while Amphiphysin, PIPK1γ and Dynamin-1 were not increased. The increase in AP-180 may reflect that some endocytic proteins can be further recruited to synapses in response to strong depolarization, matching previous reports (Bolz et al., 2023; Mueller et al., 2004).

To assess subsynaptic protein localizations, we analyzed side-view and en-face synapses after acquiring images with STED microscopy (Fig. 1 - figure supplement 2A-E). Side-view analyses enable the assessment of protein distributions axially from the plasma membrane to the inside of the bouton. We identified side-view synapses as those containing an elongated area of an active zone marker (Bassoon or Munc13) at the edge of a Synaptophysin cloud. We then extracted the distribution of the protein of interest by positioning a line profile perpendicular to the active zone marker, as we did before (Chin and Kaeser, 2024; Emperador-Melero et al., 2024, 2021b; Held et al., 2020; Tan et al., 2022). At untreated synapses, the distributions of Amphiphysin, PIPK1γ, AP-180 and Dynamin-1 were broader than those of active zone proteins (Fig. 1B-I), consistent with localization at the periactive zone and in the presynaptic cytosol (Ganguly et al., 2021; Gerth et al., 2017; Wenk et al., 2001). The average peak signal intensities of these proteins were within ∼100 nm of the peak of the active zone marker (Fig. 1J). To estimate levels at the presynaptic plasma membrane, we measured the signal intensities within a 135 nm region centered around the peak of the active zone marker (Wong et al., 2018), which likely includes the active zone and the periactive zone because the two regions overlap in side-view. The signal intensities for Amphiphysin and PIPK1γ in this region were increased upon chronic silencing, matching the higher synaptic levels quantified in confocal images, and the intensities of AP-180 and Dynamin-1 were similar to the untreated condition (Fig. 1K). The levels of AP-180, but not of the other three proteins, increased with KCl depolarization (Fig. 1K), matching the confocal data (Fig. 1 - figure supplement 1A+B). We observed a similar increase in AP-180 signals using an alternate antibody (Fig. 1 - figure supplement 3). We did not detect changes in the axial distribution of any of these proteins (Fig. 1J). Overall, these observations suggest that the localization of endocytic proteins to the presynaptic plasma membrane does not require action potential firing and presynaptic Ca^2+^ entry but instead indicate activity-independent recruitment.

Next, we analyzed en-face synapses to assess the lateral protein distribution near the plane of the plasma membrane (Fig. 1 - figure supplement 2F-I). We identified en-face synapses as those that did not have a bar-like appearance of the marker; instead, the area of the marker was surrounded by Synaptophysin staining (Emperador-Melero et al., 2024). At untreated synapses, Amphiphysin, AP-180 and Dynamin were organized into multiple clusters adjacent to the active zone. On average, these clusters were located 200 to 300 nm lateral from the center of the active zone marker (Fig. 1L-Q). The average distribution of PIPK1γ appeared closer to the active zone, possibly reflecting its roles in the synthesis of phosphatidylinositol 4,5-bisphosphate (PI(4,5)P_2_), a phospholipid important for both exo- and endocytosis (Bai et al., 2003; Bolz et al., 2023; Di Paolo et al., 2004; Wenk et al., 2001). Neither chronic silencing nor strong depolarization changed the lateral distribution of these proteins. Instead, they remained distributed in clusters surrounding the active zone (Fig. 1L-Q). There was a slight increase in the number of PIPK1γ objects upon depolarization (Fig. 1P). Upon chronic silencing, the integrated intensity of endocytic protein clusters (defined as the product of their size and average intensity) increased or showed a positive trend for all proteins (Fig. 1 - figure supplement 4). The integrated intensity also increased for AP-180 upon depolarization, matching with the higher levels measured at side-view synapses and at confocal resolution (Fig. 1 and Fig. 1 - figure supplement 4).

Together, these experiments indicate that while endocytic proteins are not impervious to synaptic activity, they are efficiently deployed to presynaptic terminals and to the periactive zone after pharmacological silencing of firing activity and Ca^2+^ entry.

### Localization of endocytic proteins after chronic silencing or acute stimulation of Drosophila NMJs

We next asked whether recruitment of endocytic proteins to the presynaptic plasma membrane follows a similar pattern at the *Drosophila* neuromuscular junction (NMJ). This is a widely used model to study synaptic architecture that contains well-defined periactive zones (Harris and Littleton, 2015). First, we used antibody labeling and STED microscopy to characterize the degree to which the endocytic proteins Dynamin, Endophilin-A (EndoA) and Dap160 (the homologue of the mammalian Intersectin) localize to the synaptic vesicle cloud or periactive zone (Fig. 2 and Fig. 2 - figure supplement 1). To do so, we measured their co-localization with the endocytic protein Nervous Wreck (FCHSD2 homologue), which exhibits strongly polarized localization to the periactive zone (Del Signore et al., 2023), or with the vesicle pool marker Synapsin. In all cases, the colocalization of these proteins with Nervous Wreck or Synapsin was partial, supporting broad distributions spanning the periactive zone and the vesicle cloud, and aligning with the distributions of endocytic proteins in mouse hippocampal neurons (Fig. 1) and with previous reports (Bai et al., 2010; Ganguly et al., 2021; Gerth et al., 2017; Wenk et al., 2001).

**Figure 2.**
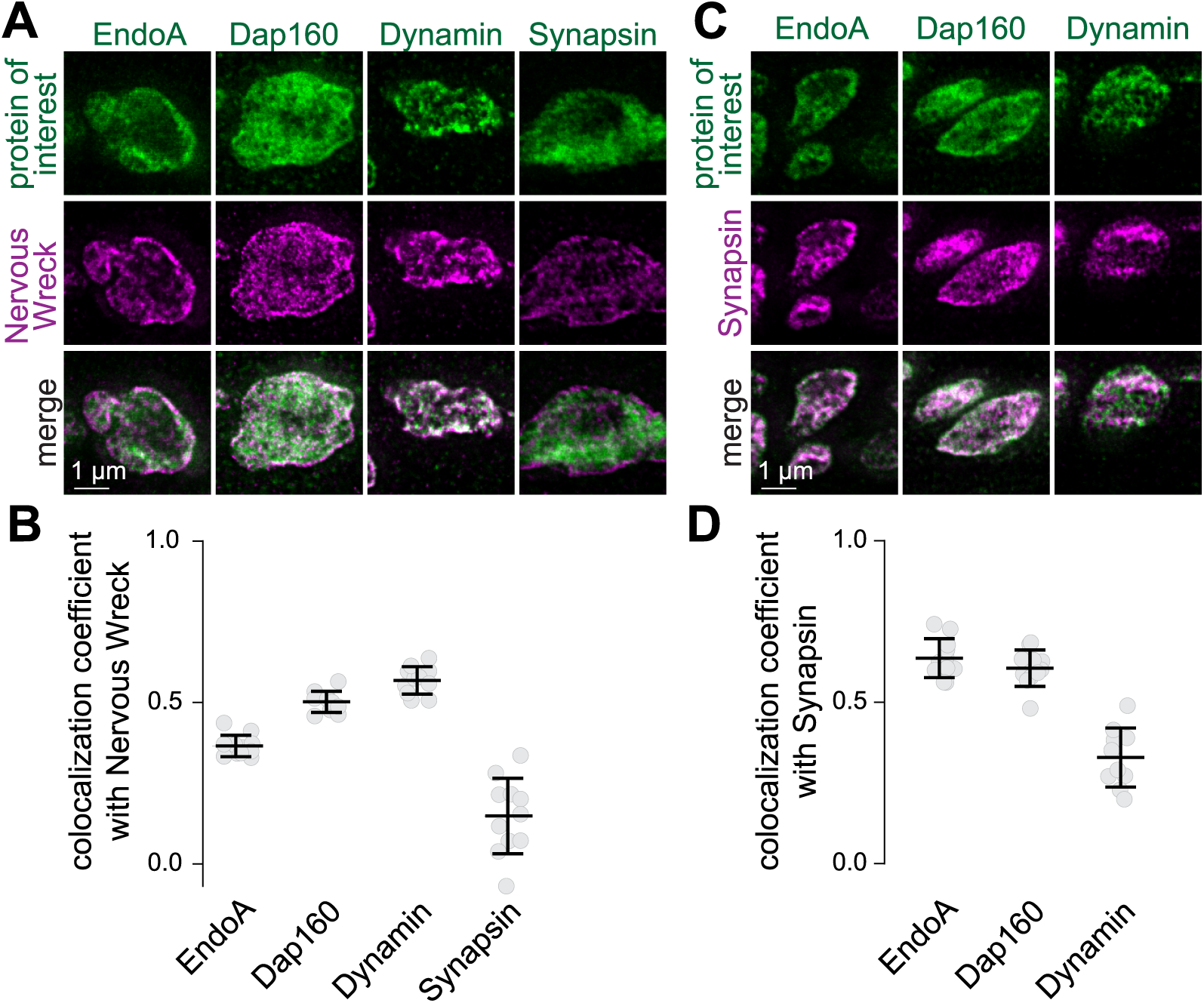
Localization of endocytic proteins at *Drosophila* NMJs relative to Synapsin or Nervous Wreck. (**A, B**) Example boutons of *Drosophila* NMJs stained for Nervous Wreck and either EndoA, Dap160, Dynamin or Synapsin, and quantification of the colocalization between Nervous Wreck and these proteins measured as the Pearson’s coefficient; n (NMJ/animals): EndoA 11/3, Dap160 10/3, Dynamin 11/3, Synapsin 11/3. (**C, D**) Same as A+B but relative to Synapsin instead of Nervous Wreck; n (NMJ/animals): EndoA 11/3, Dap160 11/3, Dynamin 10/3. Data are shown as mean ± SEM. Images acquired by STED microscopy. For antibody validation, see Fig. 2 - figure supplement 1.

Next, we assessed whether the distribution of endocytic proteins is changed by synaptic activity. We silenced Type 1b motor neurons on muscle 1 by expressing Tetanus Neurotoxin (TeNT). We employed a single-neuron driver (Jenett et al., 2012), as silencing NMJs broadly is lethal (Sweeney et al., 1995), and used motor neurons expressing the GAL4 driver alone as controls (Fig. 3a). We then used antibody staining against the active zone marker Bruchpilot (Brp), which is the homologue of ELKS (Kittel et al., 2006; Wagh et al., 2006), and combinations of the endocytic proteins Dynamin, Nervous Wreck, EndoA and Dap160, followed by Airyscan confocal microscopy and analyses of protein levels and distribution. First, we determined recruitment or retention of endocytic machinery at the terminal by measuring their mean intensity within the entire volume of the terminal. Second, to quantify deployment to the periactive zone, we used an established workflow to segment NMJs into single synaptic units composed of a periactive zone surrounding a center region containing Brp objects (Fig. 1 - figure supplement 2J-N) (Del Signore et al., 2023). Each unit is divided into ‘mesh’ and ‘core’ regions, where the periactive zone mesh is a ∼175 nm wide area localized at ∼330 nm from the center, and the ‘core’ region is the interior to this mesh (Del Signore et al., 2023). We estimated the levels of proteins in the periactive zone mesh as their mean intensity levels within this region and their distribution as the log ratio of the average intensity within the mesh over the core, with positive ratios indicating mesh enrichment and negative ratios indicating core enrichment (Fig. 1 - figure supplement 2J-N). As reported previously (Akbergenova et al., 2025), TeNT expression resulted in fewer Brp objects per µm^2^ of NMJ and an increase in their integrated intensity (Fig. 3B-D). However, TeNT expression did not alter the total levels of Nervous Wreck, Dynamin, EndoA, or Dap160, and only resulted in a small decrease of the levels of EndoA at the periactive zone mesh, and in small changes in the polarization of Dynamin and Dap160 (Fig. 3E-G). A separate experiment in which we imaged EndoA and Dap160 in these conditions using STED microscopy rendered the same conclusion (Fig. 3 - figure supplement 1), supporting the model of activity-independent recruitment of endocytic proteins at *Drosophila* NMJs.

**Figure 3.**
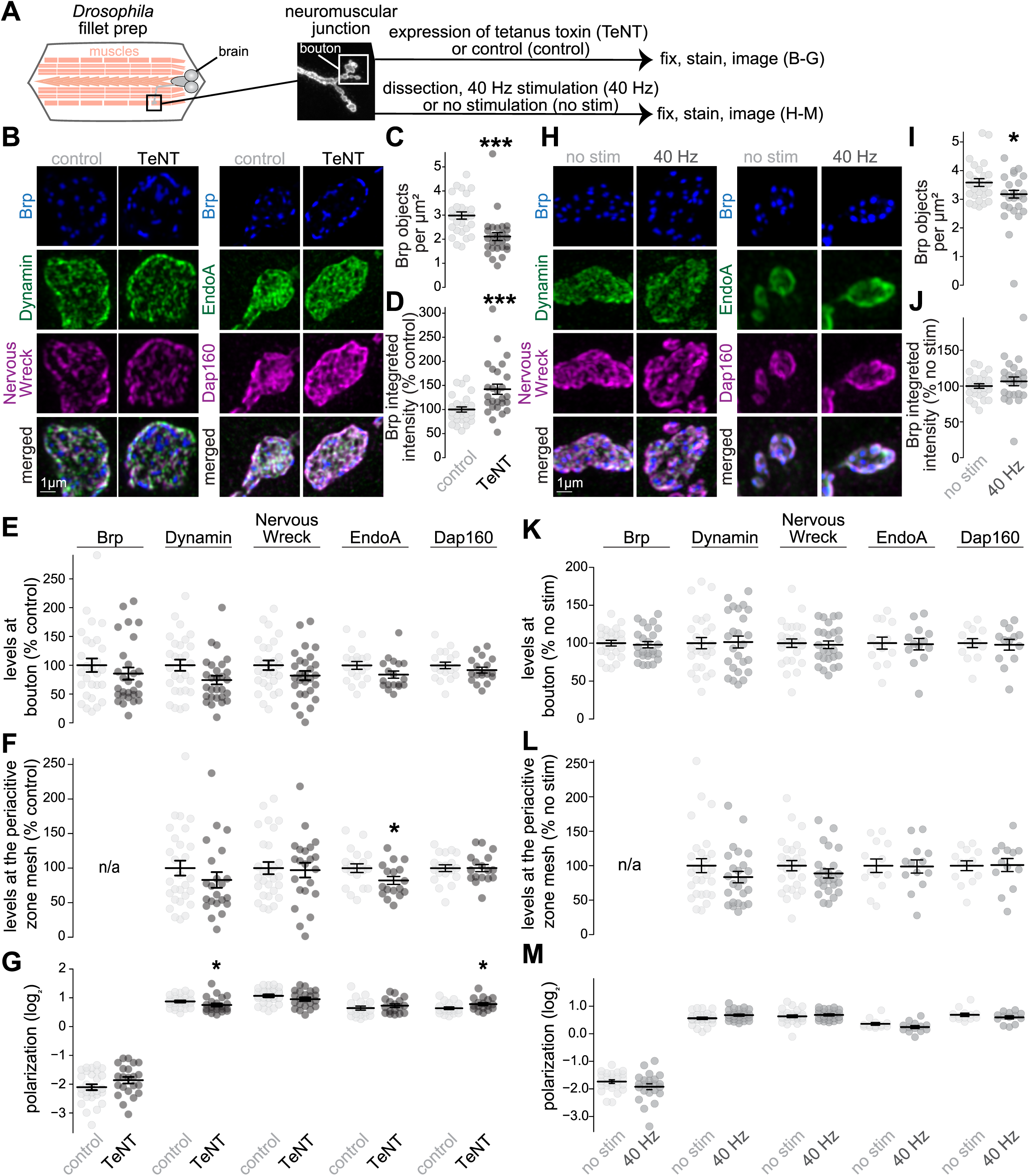
Deployment of endocytic proteins after chronic silencing or acute stimulation of *Drosophila* NMJs. (**A**) Schematic of the experiment at the *Drosophila* neuromuscular junction (NMJ). To silence neurons chronically, tetanus neurotoxin light chain (“TeNT”) was expressed in Type 1b motor neurons on muscle 1 using the GAL4 UAS system. These neurons were compared to those from larvae expressing the M11b-GAL4 driver alone (“control”). To activate neurons, electrical stimulation was applied at 40 Hz for three minutes (“40 Hz”) to Type 1b motor neurons on muscles 4 and 6/7, and comparisons were made to the contralateral, unstimulated side (“no stim”). (**B-G**) Example maximum intensity projections of ventral half of individual boutons with or without TeNT-expression stained for Brp, Dynamin and Nervous Wreck or Brp, EndoA and Dap160 (B), and quantification of the number of Brp objects per µm^2^ of NMJ (C) and their integrated intensity (D). For Brp, Dynamin, Nervous Wreck, EndoA and Dap160, the average fluorescence intensity per bouton (E), the average intensity at the periactive zone mesh (F) and the polarization within periactive zone units (G) were quantified; n in C,D (NMJs/larvae): control 28/5, TeNT 27/6; E for Nervous Wreck, Dynamin and Brp: control 28/5, TeNT 28/6; E for Dap160 and EndoA: control 19/6, TeNT 18/6; F,G for Nervous Wreck, Dynamin and Brp: control 28/5, TeNT 22/6; F,G for Dap160 and EndoA: control 19/6, TeNT 17/6. (**H-M**) Same as B-G but comparing acutely stimulated (“40 Hz”) and unstimulated terminals (“no stim”) boutons; n in I, J: no stim 26/14, 40 Hz 24/14; K for Nervous Wreck, Dynamin and Brp: no stim 26/14, 40 Hz 25/14; K for Dap160 and EndoA: no stim 13/7, 40 Hz 13/7; L,M for Nervous Wreck, Dynamin and Brp: no stim 26/14, 40 Hz 23/14; L,M for Dap160 and EndoA: no stim 13/7, 40 Hz 13/7. Data in D-F and J-L are normalized to the average of the control condition. Data are mean ± SEM; *p < 0.05 determined by two-sided Student’s t-tests (E for Nervous Wreck, EndoA and Dap160; F for EndoA and Dap160; G for Brp, Nervous Wreck, EndoA and Dap160; I-L for EndoA and Dap160; M for Brp, Dynamin and Nervous Wreck) or two-sided Mann-Whitney U tests (C-E for Brp and Dynamin; F,G for Dynamin; L,M for EndoA and Dap160). Images acquired by Airyscan microscopy, for quantification of EndoA and Dap160 using STED microscopy, see Fig. 3 - figure supplement 1.

We next tested the prediction that enhanced activity might increase the localization of endocytic proteins to the periactive zone at the *Drosophila* NMJ. To do so, we electrically stimulated Type 1b motor neurons on muscles 4 and 6/7 with a 3-minute 40 Hz train (Fig. 3A), a stimulus which induces endocytosis and causes redistribution of PI(4,5)P_2_ at the presynaptic membrane (Li et al., 2020). If deployment were enhanced by activity, we would expect increased periactive zone targeting of endocytic proteins and a concomitant increase in their polarization. However, we only observed an effect on the number of Brp objects, and saw no changes in the levels of Dynamin, Nervous Wreck, EndoA or Dap160 at boutons or at periactive zones, and no changes in the polarization of these proteins (Fig. 3H-M).

Altogether, we conclude that evoked neurotransmission is not necessary to target endocytic machinery to NMJs or to localize them within boutons, nor does enhanced activity in the tested paradigms boost their periactive zone localization.

### Efficient deployment of endocytic proteins after Ca_V_2 ablation in mouse hippocampal neurons

Action potential-triggered fusion of synaptic vesicles depends on Ca^2+^ influx via voltage-gated Ca^2+^ channels of the Ca_V_2 family (Cao et al., 2004; Cunningham et al., 2022; Held et al., 2020). As a complementary approach to study the role of synaptic activity in the recruitment of endocytic machinery, we assessed synapses lacking Ca_V_2s (Fig. 4A). We cultured neurons from previously generated triple conditional Ca_V_2 knockout mice (Ca_V_2.1, Ca_V_2.2 and Ca_V_2.3) and used lentiviral transduction to express Cre recombinase to generate Ca_V_2 triple knockout (cTKO^Cav2^) neurons. This disrupts action potential-evoked exocytosis (Chin and Kaeser, 2024; Held et al., 2020). We expressed a recombination-deficient Cre enzyme that is truncated to generate control (control^Cav2^) neurons (Held et al., 2020). In confocal microscopic images, the average synaptic signal intensities of Amphiphysin, PIPK1γ, AP-180 and Dynamin-1 were intact at cTKO^Cav2^ synapses (Fig. 4 - figure supplement 1), confirming efficient targeting of these proteins. No major changes in their subsynaptic localization were observed. When assessed in STED microscopic images, the axial protein distributions of Amphiphysin, PIPK1γ, AP-180 and Dynamin-1 were unchanged at cTKO^Cav2^ synapses, and the peak fluorescence intensities within the periactive zone region of cTKO^Cav2^ synapses were within ∼10% of the intensities measured at control^Cav2^ synapses (Fig. 4B-K and Fig. 4 - figure supplement 2). At en-face synapses, the number and intensity of clusters positioned at 100 to 300 nm from the active zone center were unchanged (Fig. 4L-Q and Fig. 4 - figure supplement 3).

**Figure 4.**
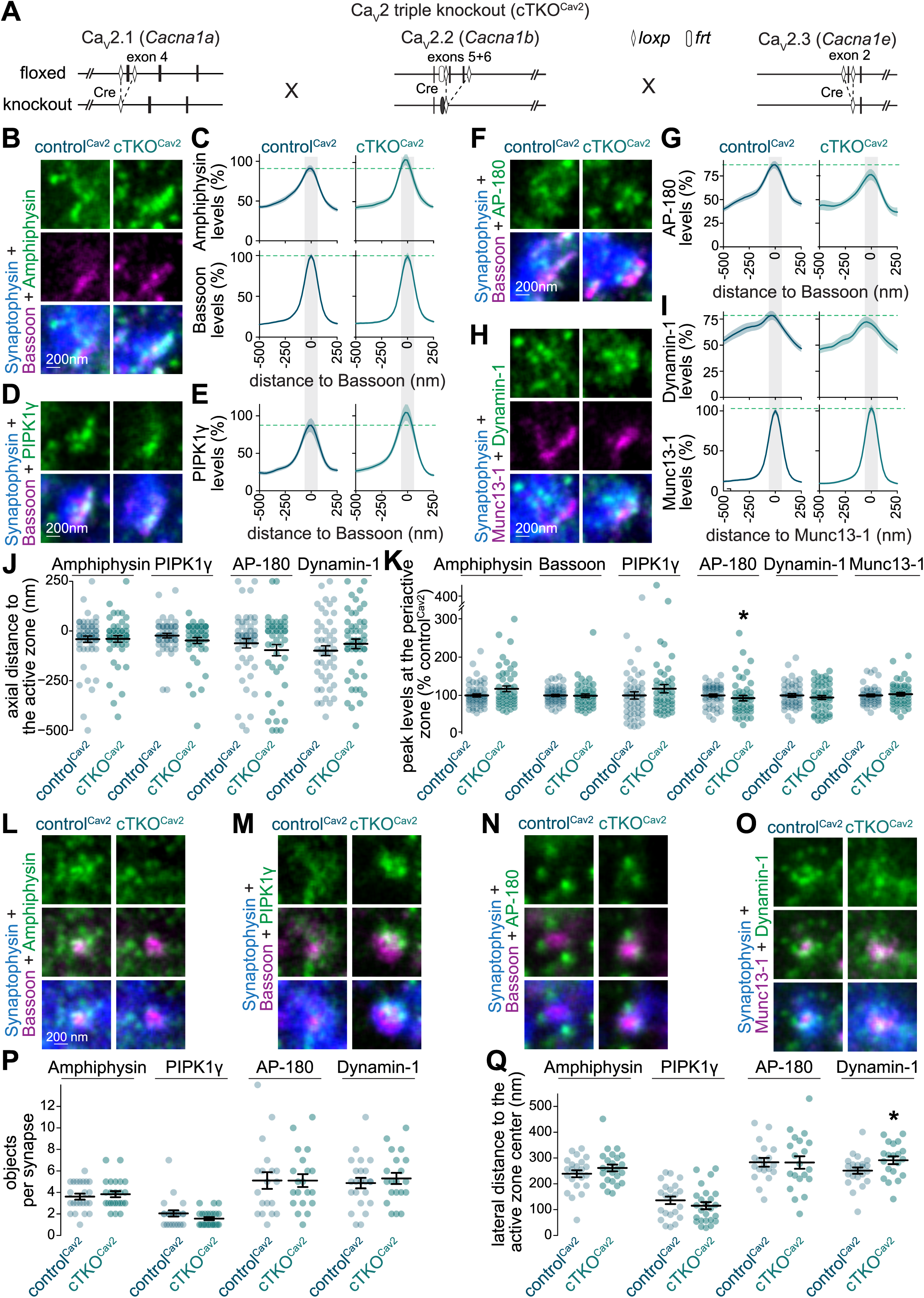
Deployment of endocytic proteins after Ca_V_2 ablation in mouse hippocampal neurons. (**A**) Schematics of the *Cacna1a*, *Cacna1b* and *Cacna1e* mutant alleles that constitute the conditional Ca_V_2 triple knockout mouse line as described in (Held et al., 2020). (**B-I**) Example side-view synapses and average line profiles of Amphiphysin and Bassoon (B,C), PIPK1γ (D,E), AP-180 (F,G), and Dynamin-1 and Munc13-1 (H,I). Neurons were stained for a protein of interest (Amphiphysin, PIPK1γ, AP-180 or Dynamin-1; imaged in STED), an active zone marker (Bassoon or Munc13-1; imaged in STED), and Synaptophysin (imaged in confocal). An area of interest was positioned perpendicular to the center of the active zone marker, and synapses were aligned via the peak fluorescence of the active zone marker in the average profiles (C, E, G, I). Line profiles plots were normalized to the average signal in the control^Cav2^ condition. Dashed lines in C, E, G and I mark average levels in the control^Cav2^ condition and grey shaded areas represent the active zone and periactive zone area; n in B, C (synapses/cultures): control^Cav2^ 58/3, cTKO^Cav2^ 50/3; D, E: control^Cav2^ 50/3, cTKO^Cav2^ 49/3; F, G: control^Cav2^ 50/3, cTKO^Cav2^ 47/3; H, I: control^Cav2^ 48/3, cTKO^Cav2^ 50/3. (**J, K**) Quantification and statistical analyses of the experiment shown in B-I, including peak-to-peak distance of the active zone marker and the protein of interest (J), and of the peak intensity in the periactive zone area (K). The periactive zone area is defined as an area within 68 nm on each side of the peak of the active zone marker (grey shaded areas in C, E, G and I); n as in B-I. (**L-Q**) Example en-face synapses (L-O) and quantification of the number of objects (P) and distance of these objects to the center of the active zone marker (Q) of the experiment shown in B-L; n in P, Q: Amphiphysin, control^Cav2^ 23/3, cTKO^Cav2^ 24/3; PIPK1γ, control^Cav2^ 22/3, cTKO^Cav2^ 25/3; AP-180, control^Cav2^ 19/3, cTKO^Cav2^ 20/3; Dynamin-1, control^Cav2^ 23/3, cTKO^Cav2^ 20/3. Data are mean ± SEM; * p < 0.5 determined by two-sided Student’s t-tests (J for Dynamin-1; K for Dynamin-1; P for PIPK1γ and Dynamin; Q for Amphiphysin, AP-180 and Dynamin-1) or two-sided Mann-Whitney U tests (J for Amphiphysin, PIPK1γ, AP-180; K for Amphiphysin, Bassoon, PIPK1γ, AP-180 and Munc13-1; P for Amphiphysin and AP-180; Q for PIPK1γ). For quantification of confocal signals, see Fig. 4 - figure supplement 1; for assessment of AP-180 using an independent antibody, see Fig. 4 - figure supplement 2; for additional assessment of en-face synapses, see Fig. 4 - figure supplement 3.

We conclude that ablating Ca_V_2s to disrupt action potential-induced synaptic vesicle exocytosis does not impair the localization of endocytic proteins, matching the outcomes of the pharmacological inhibition experiments.

### Deployment of endocytic proteins after active zone disruption in mouse hippocampal neurons

The active zone generates sites for synaptic vesicle fusion and clusters Ca_V_2 channels at the presynaptic plasma membrane (Emperador-Melero and Kaeser, 2020; Sudhof, 2012). It is a multiprotein machinery formed by RIM, RIM-BP, Liprin-α, Munc13, ELKS, Piccolo/Bassoon and other proteins that is subdivided into distinct sub-machineries (Acuna et al., 2016; Aravamudan et al., 1999; Biederer et al., 2017; Böhme et al., 2016; Emperador-Melero et al., 2024, 2021b; Emperador-Melero and Kaeser, 2020; Graf et al., 2012; Kaufmann et al., 2002; Kittel et al., 2006; Liu et al., 2011; Marcó de la Cruz et al., 2024; McDonald et al., 2020; Sudhof, 2012; Wang et al., 2016; Zhen and Jin, 1999). Endocytic machineries are localized adjacent to active zones, and the functions of these two machineries are coordinated. Hence, their assembly may be linked with instructive roles of the active zone. To test this hypothesis, we used quadruple conditional ablation of RIM1, RIM2, ELKS1 and ELKS2 at hippocampal synapses. At these synapses, active zone assembly is strongly impaired with disrupted localization of Munc13, Piccolo/Bassoon, RIM-BP and Ca_V_2 (Emperador-Melero et al., 2024; Tan et al., 2022; Wang et al., 2016; Wong et al., 2018). We cultured neurons from these mice and transduced them with lentiviruses expressing Cre or inactive Cre to produce RIM+ELKS quadruple knockout (cQKO^R+E^) or control neurons (control^R+E^), respectively (Fig. 5A). We antibody-stained these cultures as in Figs. 1 and 4, but used the postsynaptic marker PSD-95 instead of an active zone marker because the active zone is disrupted at cQKO^R+E^ synapses. At side-view synapses, PSD-95 is localized apposed to active zones at a distance of ∼75 nm (Emperador-Melero et al., 2024, 2021b; Held et al., 2020; Tan et al., 2022; Tang et al., 2016; Wong et al., 2018). Altogether, endocytic machinery remained correctly localized after active zone disruption. In confocal images (Fig. 5 - figure supplement 1), average intensities of fluorescence signals for antibody staining for Amphiphysin, PIPK1γ, AP-180 and Dynamin-1 at presynapses were either similar or higher in cQKO^R+E^ neurons. At side-view synapses in STED microscopy (Fig. 5B-L and Fig. 5 - figure supplement 2), the peak intensities were at a similar distance from PSD-95, and the signals within the region corresponding to the periactive zone were either higher or unchanged. Similarly, endocytic proteins at en-face synapses (Fig. 5M-R and Fig. 5 - figure supplement 3) from cQKO^R+E^ neurons were organized into clusters with comparable localization and similar or greater integrated density. We conclude that the localization of endocytic proteins to the periactive zone is not impaired after disrupting active zone assembly through knockout of RIM and ELKS.

**Figure 5.**
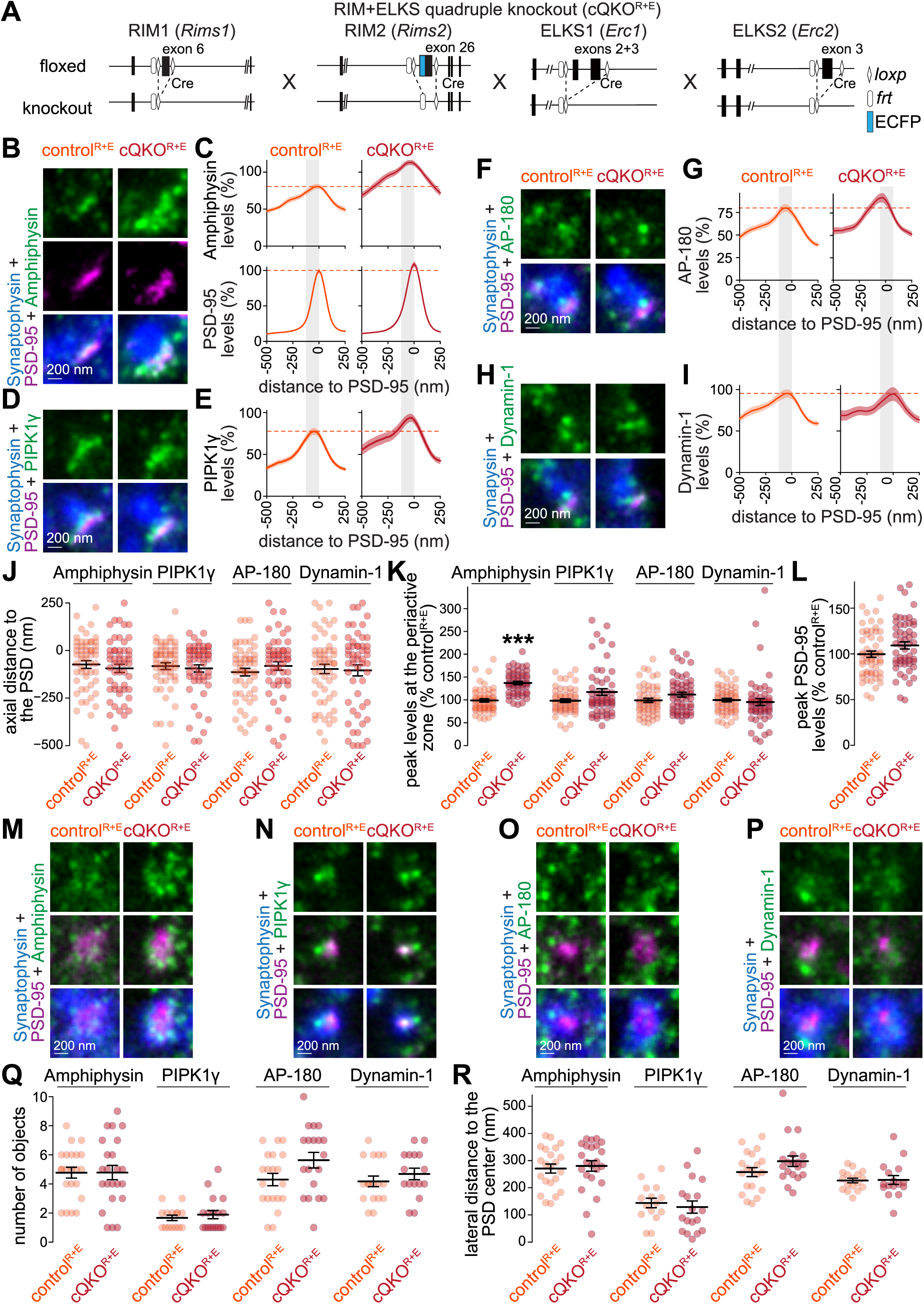
Recruitment of endocytic proteins to the periactive zone after active zone disruption in mouse hippocampal neurons. (**A**) Schematics of the *Rims1*, *Rims2*, *Erc1* and *Erc2* mutant alleles that constitute the conditional RIM+ELKS quadruple knockout mouse line as described in (Tan et al., 2022; Wang et al., 2016). (**B-I**) Example side-view synapses and average line profiles of Amphiphysin and PSD-95 (B,C), PIPK1γ (D,E), AP-180 (F,G), and Dynamin-1 (H,I). Neurons were stained for a protein of interest (Amphiphysin, PIPK1γ, AP-180 or Dynamin-1; imaged in STED), a postsynaptic marker (PSD-95; imaged in STED), and a synaptic vesicle marker (Synaptophysin or Synapsin; imaged in confocal). An area of interest was positioned perpendicular to the center of the PSD-95 object, and synapses were aligned via the peak fluorescence of PSD-95 in the average line profiles (C, E, G, I). Line profile plots were normalized to the average signal in the control^R+E^ condition. Dashed lines in C, E, G and I mark average levels in the control^R+E^ condition and grey shaded areas represent the active zone and periactive zone area; n in B, C (synapses/cultures): control^R+E^ 55/3, cQKO^R+E^ 53/3; D, E: control^R+E^ 55/3, cQKO^R+E^ 54/3; F, G: control^R+E^ 53/3, cQKO^R+E^ 54/3; H, I: control^R+E^ 54/3, cQKO^R+E^ 53/3. (**J-L**) Quantification and statistical analyses of the experiment shown in B-I, including-to-peak distance of PSD-95 and the protein of interest (J), peak intensity in the periactive zone area (K), and peak intensity of PSD-95 (L). The periactive zone area is defined as an area between -136 nm and the peak of PSD-95 (grey shaded areas in C, E, G and I); n as in B-I. (**M-R**) Example en-face synapses (M-P) and quantification of the number of objects (Q) and distance of these objects to the center of the PSD-95 object (R) of the experiment shown in B-L; n in Q, R: Amphiphysin, control^R+E^ 22/3, cQKO^R+E^ 23/3; PIPK1γ control^R+E^ 15/3, cQKO^R+E^ 18/3; AP-180 control^R+E^ 20/3, cQKO^R+E^ 19/3; Dynamin-1 control^R+E^ 17/3, cQKO^R+E^ 16/3. Data are mean ± SEM; ***p < 0.001 as determined by two-sided Student’s t-tests (L,Q for Amphiphysin, AP-180 and Dynamin-1; R for PIPK1γ and Dynamin-1) or two-sided Mann-Whitney U tests (J, K, Q for PIPK1γ; R for Amphiphysin and, AP-180). For quantification of confocal signals, see Fig. 5 - figure supplement 1; for assessment of AP-180 using and independent antibody, see Fig. 5 - figure supplement 2; for additional assessment of en-face synapses, see Fig. 5 - figure supplement 3.

### Deployment of endocytic proteins at Drosophila NMJs after disrupting active zone assembly

The active zone of the *Drosophila* NMJ has a T-bar, a prominent electron-dense invagination consisting mainly of Brp (Fouquet et al., 2009; Kittel et al., 2006; Wagh et al., 2006). Brp is required for the normal clustering of Ca_V_2 Ca^2+^ channels at the active zone, and its ablation severely decreases evoked release (Fouquet et al., 2009; Kittel et al., 2006; Wagh et al., 2006). To determine whether these defects perturb periactive zone protein recruitment, we analyzed levels and localization of endocytic proteins in *brp* mutants (Fig. 6A). After genetic ablation of *brp*, the levels of Dynamin and Nervous Wreck at NMJs were not decreased and these proteins remained concentrated in the periactive zone mesh (Fig. 6B-E). These data indicate that Brp and active zone T-bars are dispensable for the localization of these endocytic proteins to the periactive zone at the *Drosophila* NMJ and support the finding (Fig. 1B-G and 3B-G) that periactive zone properties do not depend on evoked release.

**Figure 6.**
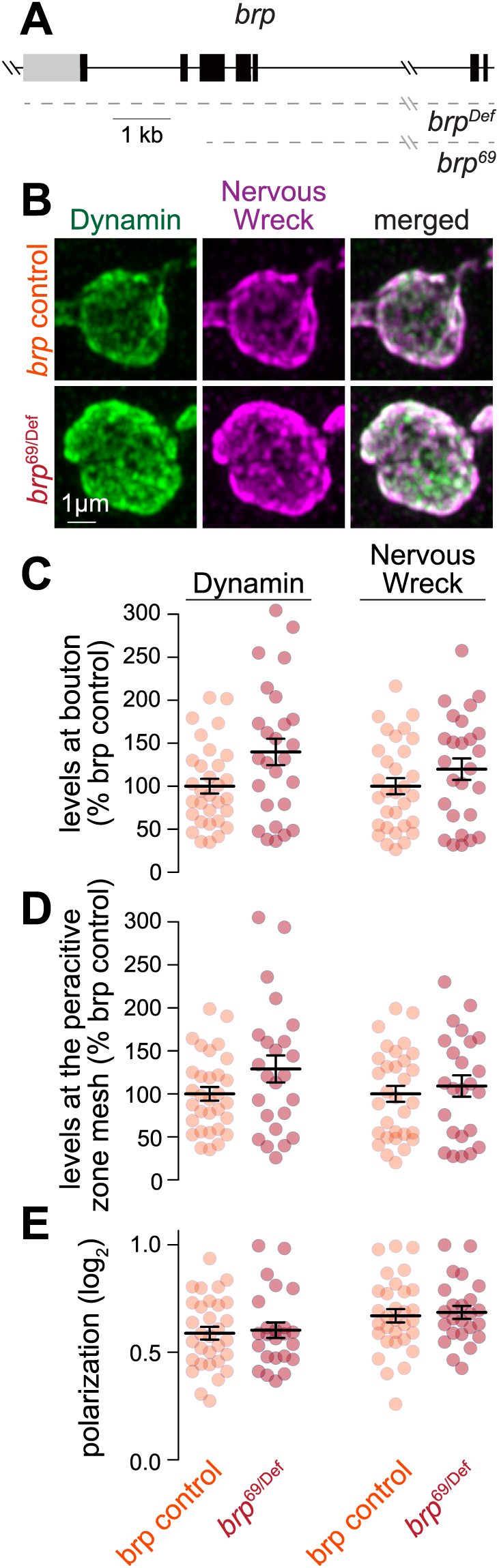
Recruitment of endocytic proteins to the periactive zone after disrupting active zone assembly at *Drosophila* NMJs. (**A**) Schematic of the *brp^69^* and *brp^Def^* alleles; dashed lines indicate the extent of deletions. (**B-E**) Example maximum intensity projections of individual boutons of muscle 4 Type 1b terminals from control *white* and *brp^69^/brp^Def^* larvae (B) and quantification of the average fluorescence intensity per bouton (C), average intensity at the periactive zone mesh (D) and polarization within periactive zone units (E) for Dynamin and Nervous Wreck. Data in C and D are normalized to the average control condition; n in C (NMJs/larvae): *brp* control 31/3, *brp^69/Def^* 26/3; D+E, *brp* control 31/3, *brp^69/Def^* 24/3. Data are mean ± SEM; statistical significance assessed using two-sided Student’s t-tests (C for Nervous Wreck; D for Nervous Wreck; E for Nervous Wreck) or two-sided Mann-Whitney U tests (C for Dynamin; D for Dynamin; E for Dynamin). Images acquired by Airyscan microscopy.

### Deployment of endocytic machinery after disruption of upstream active zone assembly mechanisms

The observation that active zone disassembly does not reduce levels or disrupt localization of endocytic proteins at periactive zones (Figs. 5 and 6) suggests that active zones and endocytic machineries are organized independently of one another. Alternatively, shared upstream assembly pathways might help co-organization of the active zone and the periactive zone. Liprin-α is an upstream active zone organizer that maintains priming machineries and recruits additional presynaptic material (Böhme et al., 2016; Emperador-Melero et al., 2024, 2021b; Kaufmann et al., 2002; Marcó de la Cruz et al., 2024; McDonald et al., 2020; Wong et al., 2018; Zhen and Jin, 1999). We assessed the localization of endocytic proteins in Liprin-α mutants to assess whether Liprin-α might co-organize active zone and periactive zone assembly. In vertebrates, four Liprin-α proteins are expressed from four different genes (Zurner and Schoch, 2009). We previously generated quadruple Liprin-α mutant mice containing conditional Liprin-α1, -α2 and -α4 knockout alleles and constitutive Liprin-α3 knockout alleles (Emperador-Melero et al., 2024, 2021b; Wong et al., 2018). We cultured hippocampal neurons from these mice and transduced them with Cre lentivirus to produce quadruple knockout (cQKO^L1-L4^) neurons, or a combination of two viruses to express Liprin-α3 and a truncated, inactive version of Cre, to generate control (control^L1-L4^) neurons, as described before (Emperador-Melero et al., 2024). Synapses of cQKO^L1-L4^ neurons have altered active zone properties, with decreased levels of RIM and Munc13-1, accompanied by an impairment in the readily releasable pool of synaptic vesicles (Emperador-Melero et al., 2024). Endocytic proteins were overall efficiently deployed to presynaptic terminals and correctly localized to the periactive zone in cQKO^L1-L4^ neurons (Fig. 7 and Fig. 7 - figure supplements 1 and 2). There were only small decreases in signal intensities generated by Amphiphysin and AP-180 antibodies, a small increase in PIPK1y, and a modest shift in the lateral position of Amphiphysin. These results indicate that endocytic proteins are overall efficiently localized in the absence of Liprin-α proteins in mouse hippocampal neurons.

**Figure 7.**
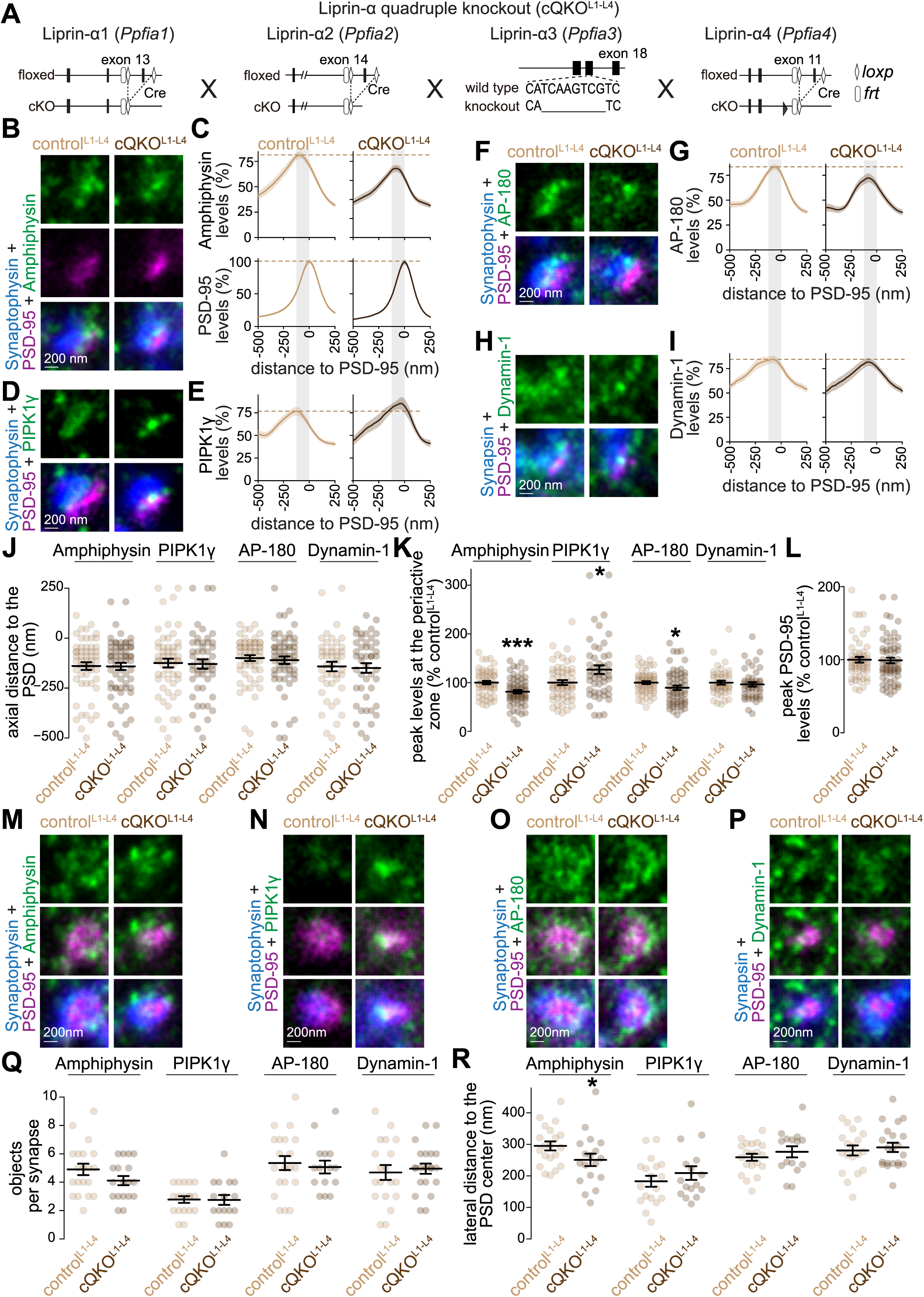
Deployment of endocytic proteins after Liprin-α ablation in mouse hippocampal neurons. (**A**) Schematics of the *Ppfia1*, *Ppfia2*, *Ppfia3* and *Ppfia4* mutant alleles that constitute the conditional Liprin-α quadruple knockout mouse line as described in (Emperador-Melero et al., 2024). (**B-I**) Example side-view synapses and average line profiles of Amphiphysin and PSD-95 (B+C), PIPK1γ (D+E), AP-180 (F+G), and Dynamin-1 (H+I). Neurons were stained for a protein of interest (Amphiphysin, PIPK1γ, AP-180 or Dynamin-1; imaged in STED), a postsynaptic marker (PSD-95; imaged in STED), and a synaptic vesicle marker (Synaptophysin or Synapsin; imaged in confocal). An area of interest was positioned perpendicular to the center of the PSD-95 object, and synapses were aligned via the peak fluorescence of PSD-95 in the average profiles (C, E, G, I). Line profile plots were normalized to the average signal in the control^L1-L4^ condition. Dashed lines in C, E, G and I mark average levels in the control^L1-L4^ condition and grey shaded areas represent the active zone and periactive zone area; n in B (synapses/cultures), C: control^L1-4^ 54/3, cQKO^L1-4^ 64/3; D, E: control^L1-4^ 55/3, cQKO^L1-4^ 50/3; F, G: control^L1-4^ 59/3, cQKO^L1-4^ 59/3; H, I: control^L1-4^ 45/3, cQKO^L1-4^ 46/3. (**J-L**) Quantification and statistical analyses of the experiment shown in B-I, including peak-to-peak distance of PSD-95 and the protein of interest (J), peak intensity in the periactive zone area (K), and peak intensity of PSD-95 (L). The periactive zone area is defined as an area between -136 nm and the peak of PSD-95 (grey shaded areas in C, E, G and I); n as in B-I. (**M-R**) Example en-face synapses (M-P) and quantification of the number of objects (Q) and distance of these objects to the center of the PSD-95 object (R) of the experiment shown in B-L; n in Q, R: Amphiphysin, control^L1-4^ 20/3, cQKO^L1-4^ 18/3; PIPK1γ control^L1-4^ 18/3, cQKO^L1-4^ 16/3; AP-180 control^L1-4^ 20/3, cQKO^L1-4^ 15/3; Dynamin-1 control^L1-4^19/3, cQKO^L1-4^ 21/3. Data are mean ± SEM; *p < 0.05, ***p < 0.001 determined by two-sided Student’s t-tests (J for PIPK1γ; K for Amphiphysin; Q for Amphiphysin, AP-180 and Dynamin-1, R for Amphiphysin, AP-180 and Dynamin-1) or two-sided Mann-Whitney U tests (J for Amphiphysin, AP-180 and Dynamin-1; K for PIPK1γ, AP-180 and Dynamin-1; L,Q for PIPK1γ; R for PIPK1γ). For quantification of confocal signals, see Fig. 7 - figure supplement 1; for additional assessment of en-face synapses, see Fig. 7 - figure supplement 2.

To disrupt *liprin-α* in *Drosophila*, we used a previously characterized heteroallelic mutant that results in loss-of-function of this protein (*liprin-α*^EPexR60^/*liprin-α*^F3ex15^) (Fouquet et al., 2009; Kaufmann et al., 2002; Owald et al., 2010) (Fig. 8A). Loss of *liprin-α* function reduced the overall levels of Brp at NMJs and resulted in fewer Brp puncta with a greater integrated intensity (Fig. 8B-E) as described previously (Fouquet et al., 2009; Kaufmann et al., 2002; Owald et al., 2010)). In contrast, the bouton and periactive zone levels of Dynamin, Nervous Wreck, EndoA and Dap160 remained similar between controls and *liprin-α* mutants, though we did detect small-magnitude decreases in the polarization of Dynamin, Nervous Wreck and Dap160 (Fig. 8E-G). We conclude that Liprin-α does not function as a major organizer of endocytic machinery at *Drosophila* NMJs, a finding that further highlights the independence between active zone and periactive zone assembly.

**Figure 8.**
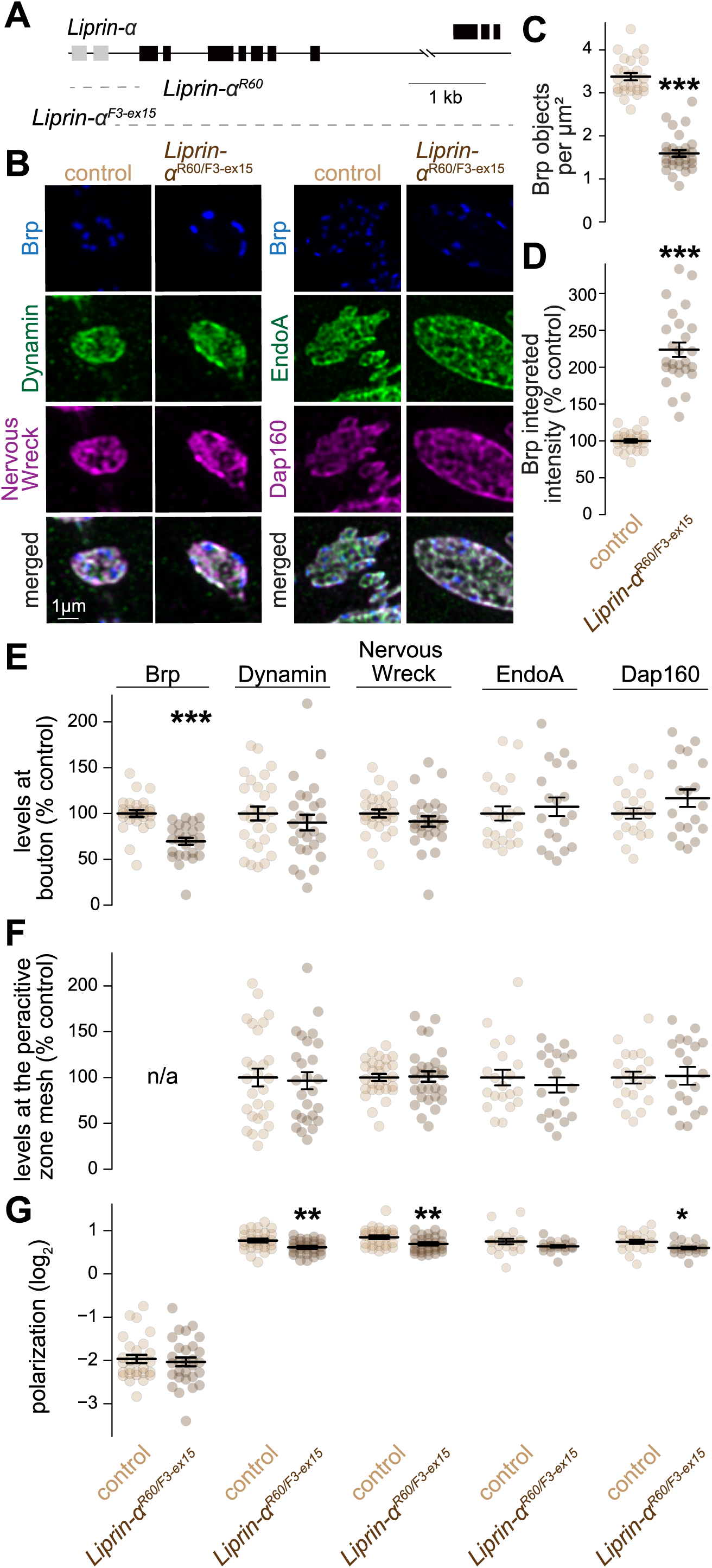
Endocytic proteins are recruited to periactive zones of *Drosophila* NMJs in *Liprin-α* mutants. (**A**) Schematic of the *Liprin-α^R60^* and *Liprin-α^F3-ex15^* alleles; dashed lines indicate the extent of deletions. (**B-G**) Example maximum intensity projections of muscle 4 Type 1b terminals from *Liprin-α^R60/F3-ex15^* mutant and *white* control larvae (B), and quantification of the number of Brp objects per µm^2^ of NMJ (C) and of their integrated intensity (D). For Brp, Dynamin, Nervous Wreck, EndoA and Dap160, the average fluorescence intensity per bouton (E), the average intensity at the periactive zone mesh (F) and the polarization within periactive zone units (G) were quantified. Data in D, E and F are normalized to the average of the control condition; n in C, D and E-G for Nervous Wreck, Dynamin and Brp (NMJs/larvae): control 27/5, *Liprin-α^R60/F3-ex15^* 26/5; E for Dap160 and EndoA: control 21/6, *Liprin-α^R60/F3-ex15^* 19/6; F,G for Dap160 and EndoA: control 20/6, *Liprin-α^R60/F3-ex15^* 19/6. Data are mean ± SEM; * p < 0.05, ** p < 0.01, *** p < 0.001 determined by two-sided Student’s t-tests (E for Dynamin and Nervous Wreck; F,G for Dynamin, EndoA and Dap160) or two-sided Mann-Whitney U tests (C-E for Brp, EndoA and Dap160; G for Brp and Nervous Wreck). Images acquired by Airyscan microscopy.

As an alternative strategy to address the independence between these two machineries at the fly NMJ, we assessed the role of the synaptic vesicle-associated protein Rab3 (Fig. 9A) using an established loss-of-function mutant (*rab3^rup^*) (Graf et al., 2009; Peled and Isacoff, 2011). *rab3* mutants had fewer but larger Brp objects (Fig. 9B-D) (Graf et al., 2009; Peled and Isacoff, 2011), though by a mechanism distinct from TeNT expression (Akbergenova et al., 2025). Neither the levels nor the distribution of Dynamin or Nervous Wreck were affected in *rab3* mutants (Fig. 9E-G), showing that their periactive zone targeting is independent of *rab3*. Notably, the localization of these endocytic proteins was similar between Brp-positive and Brp-negative periactive zones in both controls and *rab3* mutants, with only modest shifts in the polarization of Dynamin in controls and of Nervous Wreck in *rab3* mutants (Fig. 9H-K). Given the reduction in the number of Brp objects, the percentage of periactive zones without detectable Brp objects in *rab3* mutants was greater (Fig. 9L-N). Importantly, these Brp-absent periactive zones remained apposed to the postsynaptic marker Pak (Fig. 9N), arguing that Brp-negative periactive zones are not simply regions of membrane between adjacent periactive zones. We conclude that the recruitment of endocytic machinery does not require *rab3* and that periactive zones can be formed without an active zone.

**Figure 9.**
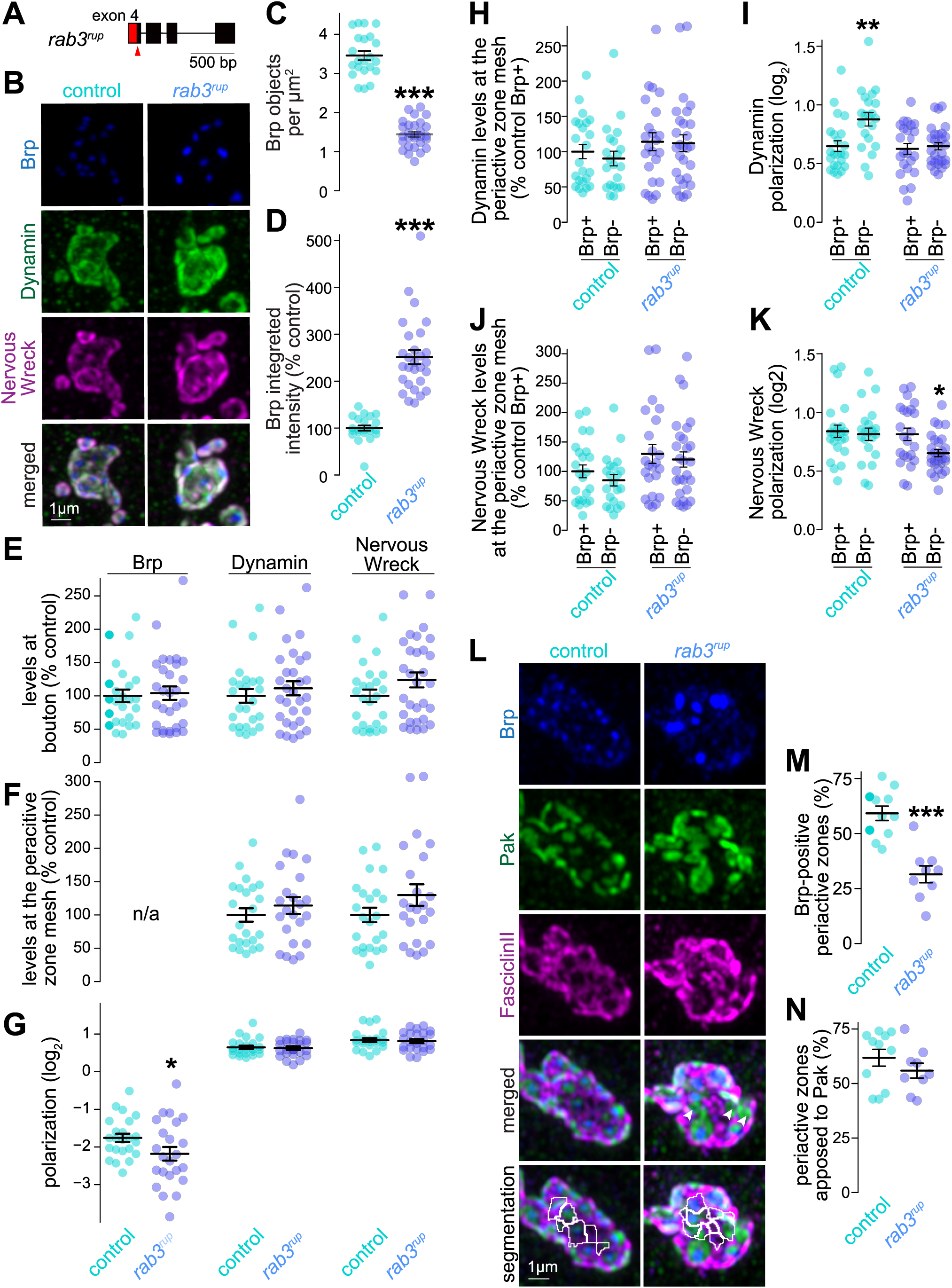
Recruitment of endocytic proteins at *Drosophila* NMJs in *rab3* mutants. (**A**) Schematic of the *rab3*^rup^ allele; red triangle indicates site of 5 bp deletion and frameshift that eliminates the final 35 amino acids (highlighted in red). (**B-G**) Example maximum intensity projections of muscle 4 Type 1b terminals from *rab3*^rup^ mutants and *white* larvae controls (B), and quantification of the number of Brp objects per µm^2^ of NMJ (C) and of their integrated intensity (D). For Brp, Dynamin and Nervous Wreck, the average fluorescence intensity per bouton (E), the average intensity at the periactive zone mesh (F) and the polarization within periactive zone units (G) were quantified. Data in D, E and F are normalized to the average of the control condition; n in C, D (NMJs/larvae): control 22/5, *rab3*^rup^ 28/6; E: control 25/5, *rab3*^rup^ 30/6; F, G: control 23/5, *rab3*^rup^ 23/6. (**H, I**) Quantification of the levels of Dynamin (H) and of its polarization (I); n in H, I: control Brp+, 23/5, control Brp- 21/6, *rab3^rup^* Brp+ 23/6, *rab3*^rup^ Brp- 28/6; (**J, K**) Same as H,I, but for Nervous Wreck; n as in H, I. (**L-N**) Example maximum intensity projections (L) of muscle 4 Type 1b terminals co-stained for the active-zone marker Brp, the postsynaptic density marker Pak, and the periactive zone marker Fasciclin-II (including the periactive zone segmentation, bottom image), and quantification of the percentage of individual periactive zone segments that contain Brp (M) or are apposed to Pak (N). Arrowheads point at periactive zones lacking Brp. n control 11/3, *rab3*^rup^ 9/3. Data are mean ± SEM; * p < 0.05, ** p < 0.01, *** p < 0.001 determined by two-sided Mann-Whitney U tests (C, D, E, F, G, M, N) or by Kruskal-Wallis tests followed by Holm post hoc tests (H-K). In H-K, data are compared to the control Brp+ condition. Images acquired by Airyscan microscopy.

Together, these data establish that the pathways organizing endocytic machinery are different from those organizing active zones.

## Discussion

We tested the hypotheses that synaptic activity or active zone scaffolds recruit and organize endocytic machinery at the periactive zone. In mouse hippocampal neurons, neither block of action potentials and presynaptic Ca^2+^ entry nor ablation of Ca_V_2s disrupted the periactive zone localization of the endocytic proteins Amphiphysin, Dynamin-1, PIPK1γ or AP-180. Similarly, at the *Drosophila* NMJ, inhibiting evoked exocytosis did not change the levels or the subsynaptic distribution of Dynamin, Nervous Wreck, EndoA or Dap160. These endocytic proteins also remained clustered at the periactive zone in both types of synapses when active zone assembly was disrupted via ablation of the scaffolds Brp or RIM and ELKS, or of Liprin-α or Rab3. Our data collectively argue that a periactive zone protein network, consisting of a high concentration of membrane-localized endocytic proteins, forms and is maintained independent of action potential-induced synaptic activity and active zone protein machinery.

### Roles for activity in endocytic assemblies

We show that a significant fraction of endocytic proteins is already deployed to the membrane prior to stimulation. Previous experiments tested the sufficiency of depolarization to induce recruitment of endocytic machinery to the periactive zone, but not whether it is necessary (Koh et al., 2007; Winther et al., 2015, 2013). We found that neither continuous silencing of neurons nor ablating Ca_V_2 channels resulted in decreased levels of endocytic proteins at the periactive zone. Instead, the levels of some of these proteins were increased in these conditions, the opposite of the effect expected for activity-dependent recruitment. A mechanism for this effect might be a homeostatic response (Wen and Turrigiano, 2024) similar in magnitude to the increase in active zone protein levels following activity blockade (Held et al., 2020). Increased synaptic enrichment was also observed for Endophilin at nematode NMJs in mutants with disrupted exocytosis (Bai et al., 2010). We do not see such large shifts in Endophilin following similar manipulations, which might reflect distinct synaptic architectures in the *C. elegans* dorsal cord vs *Drosophila* NMJ terminals.

Endocytic protein recruitment and localization were largely insensitive to acute stimulation. At mouse hippocampal synapses, Dynamin-1, Amphiphysin and PIPK1γ did not increase upon depolarization with KCl, nor did Dynamin, Nervous Wreck, EndoA or Dap160 upon electrical stimulation of *Drosophila* NMJs. Importantly, there were varying degrees of periactive zone enrichment of endocytic proteins (Fig. 2), and the data do not exclude the possibility that a subset of endocytic proteins is responsive to activity. In line with this, acute depolarization increased the clustering of AP-180 at periactive zones (Fig. 1), resembling previously reported increases in Clathrin using a similar experimental paradigm (Mueller et al., 2004), suggesting that Clathrin and associated adapters are responsive to activity. Like the active zone (Emperador-Melero et al., 2024), the endocytic machinery may be formed by distinct submachineries (Kaksonen and Roux, 2018), which may have different degrees of constitutive deployment to the periactive zone and sensitivities to synaptic activity.

### Pathways for coordinating active zone and periactive zone assembly

Despite the functional coupling of exocytosis and endocytosis in nerve terminals, it has remained unknown whether active zone proteins mediate the formation of the periactive zone. We find that ablating active zone scaffolds, upstream organizers, or Ca^2+^ channels did not disrupt the formation or structure of the periactive zone. What might be the pathways to coordinate the assembly of these two adjacent machineries? One possibility is that the co-organization of these machineries is instructed by a network of interconnected cell adhesion proteins. LAR-RPTPs and Neurexins are localized to and organize the active zone (Emperador-Melero et al., 2024, 2021a; Sclip and Südhof, 2023, 2020; Trotter et al., 2019). Cell adhesion proteins localized at the edge of a synapse, such as SynCAM or Fasciclin-II (Jiao et al., 2010; Perez de Arce et al., 2015), may play a similar role for endocytic machinery. Presynaptic cell-adhesion complexes, for example LAR-RPTPs and Neurexins, might be interlinked for the coordination of their intracellular interactors (Thivaios et al., 2024). Presynaptic protein complexes might also be organized via transcellular interactions, a model that is supported by our observation that periactive zones lacking Brp remain aligned with the postsynaptic marker Pak, pointing to trans-synaptic mechanisms for periactive zone organization.

Two additional contributing factors are the cytoskeleton and lipids. Multiple endocytic proteins bind to and function in the nucleation and polymerization of actin filaments (Del Signore et al., 2021; Saheki and De Camilli, 2012; Stanishneva-Konovalova et al., 2016). Interactions between several active zone proteins, such as Piccolo/Bassoon or Liprin-α, and actin regulators have also been described (Brenig et al., 2015; Terry-Lorenzo et al., 2016) and these interactions may be important during development for the assembly of presynaptic compartments (Chia et al., 2012). This may explain the small decrease in the levels of Amphiphysin and AP-180 that we observed in Liprin-α null neurons. Similarly, PI(4,5)P_2_ binds to RIM and this interaction is critical for synaptic strength (de Jong et al., 2018), and the synthesis of this lipid is important for endocytosis (Di Paolo et al., 2004; Wenk et al., 2001). Hence, actin and/or lipids may act as hubs around which active zone and endocytic machineries are organized.

Finally, interactions between endocytic proteins may further contribute to the anchoring of this apparatus. Most of these interactions are weak and transient, which may account for the high degree of turnover or mobility of these proteins at synapses (Reshetniak et al., 2020). However, their high concentration and their numerous interactions may suffice to maintain a stable periactive zone structure (Saheki and De Camilli, 2012; Wilhelm et al., 2014). In support of this point, perturbing interactions between Dynamin-1 and Endophilin-A1 increases the distance between these proteins (Imoto et al., 2024), suggesting their binding has a scaffolding function. Ultimately, it is plausible that several or all of these pathways organize the endocytic apparatus in parallel. Redundancy is a guiding principle in the assembly of the active zone (Acuna et al., 2016; Wang et al., 2016) and for synaptic cell adhesion (Sclip and Südhof, 2023), and it may similarly guide endocytic assemblies. Future studies should use loss-of-function and gene ablation approaches, including for Dynamins and other endocytic proteins to assess roles of these pathways in presynaptic assembly.

### Importance of the constitutively deployed endocytic apparatus for synapse function

The constitutive deployment of endocytic machinery to periactive zones likely enables multiple synaptic adaptations and functions of the endocytic machinery. First, ultrafast endocytosis, which occurs immediately following release (Watanabe et al., 2014), depends on the pre-deployment of Dynamin-1 and assembly of a periactive zone-associated ring of F-actin to facilitate membrane compression upon exocytosis (Imoto et al., 2022; Ogunmowo et al., 2023). Pre-deployment may also facilitate vesicle recycling by limiting diffusion of synaptic vesicle cargoes (Gimber et al., 2015), and may be important to prevent synaptic depression (Hua et al., 2011; Kawasaki et al., 2000; Moro et al., 2021). Finally, proteins involved in endocytosis may contribute to additional functions, such as fusion of dense core vesicles, regulation of the fusion pore, and trafficking of receptors, extracellular vesicles and cell adhesion proteins (Anantharam et al., 2011; Bailey et al., 1992; Blanchette et al., 2022; Fu and Huang, 2010; Moro et al., 2021; Rodal et al., 2008). Thus, constitutive deployment of the endocytic machinery is likely an essential adaptation to meet the demands for fast, scalable, and robust membrane internalization during synaptic vesicle recycling and to facilitate its diverse functions at the synapse.

Given the constitutive deployment of endocytic machinery, why is it not constitutively active and continuously retrieving presynaptic plasma membrane? Plausible mechanisms may include one or more of the following factors, which might be relieved in response to Ca^2+^ entry or synaptic vesicle fusion. First, the endocytic machinery may be maintained in an inactivated state via autoinhibition of some of its components, for example Endophilin or Nervous Wreck (Del Signore et al., 2021; Rao et al., 2010; Stanishneva-Konovalova et al., 2016). Second, endocytic proteins deployed to the periactive zone might be segregated into distinct periactive zone protein pools and thus not be able to functionally interact. In support of this model, we observed that the clustering pattern of PIPK1γ in silenced hippocampal synapses was different from that of Dynamin-1, Amphiphysin and AP-180 (Fig. 1). Furthermore, our previous work revealed low colocalization between Dynamin, Nervous Wreck, Dap-160 and Clathrin within periactive zones at the *Drosophila* NMJs (Del Signore et al., 2023). Third, one or multiple essential components of the endocytic machinery might not be predeployed to the membrane and may be delivered by an alternative mechanism (Bai et al., 2010). This is supported by the observation that synaptic vesicle proteins, such as Synaptotagmin-1, have a role in endocytosis (Bolz et al., 2023; Haucke and De Camilli, 1999; Jorgensen et al., 1995; Poskanzer et al., 2006; Zhang et al., 1994). Finally, the endocytic process may rely on a decrease in membrane tension (Ogunmowo et al., 2023; Watanabe et al., 2013) and thus require fusion for its initiation. Given the multiple mechanisms by which periactive zone proteins likely contribute to synaptic vesicle endocytosis, and considering their potential synaptic vesicle-independent functions, it is possible that constitutive deployment is a necessary synaptic adaptation of the conserved endocytic machinery. Testing whether this is the case will require identifying the mechanisms that are essential for constitutive deployment of the endocytic machinery to the periactive zone.

### Limitations and outlook

The presented data indicate that evoked neurotransmission and active zone scaffolds are not required for the localization of the tested endocytic proteins to the periactive zone, but several limitations are present.

First, conclusions that can be drawn on the roles of spontaneous release in periactive zone assembly remain limited. While many of the manipulations used here, including Ca_V_2 knockout (Held et al., 2020), RIM+ELKS knockout (Tan et al., 2022; Wang et al., 2016) and Liprin-α knockout (Emperador-Melero et al., 2024) in hippocampal neurons, and TeNT expression in fly NMJs (Sweeney et al., 1995), result in 50% to 70% decreased spontaneous release rates, it is possible that the remaining spontaneous release supports periactive zone assembly. Future studies might test manipulations with strong effects on miniature release including those affecting SNARE proteins and their regulators, with the caveat that these manipulations might have effects on upstream trafficking and in some cases on cell survival (Kaeser and Regehr, 2014; Santos et al., 2017).

Second, the endocytic machinery might be sensitive to manipulations over timescales that were not included in the presented analyses. The experiments with activity induction focused on periactive zone protein enrichment immediately following stimulation and do not exclude that activity may recruit endocytic proteins over slower time scales. Likewise, our data cannot exclude localization of endocytic proteins to other axonal compartments, where they may execute functions different from endocytosis.

Finally, the studies presented here are focused on the recruitment and organization of endocytic machinery, and it was not tested whether these manipulations perturb endocytic function. Studying these deficits in mutants that alter exocytosis is complicated by the fact that synaptic endocytosis occurs following exocytosis. Furthermore, functional roles for endocytic proteins beyond endocytosis, such as those for Intersectin in vesicle clustering (Milovanovic et al., 2018), and of Endophilin in autophagy (Bademosi et al., 2023; Soukup and Verstreken, 2017), may also be present.

Overall, the data on endocytic protein localization argue for constitutive deployment of this protein machinery to the periactive zone This work builds a foundation to assess alternative mechanisms and models of periactive zone assembly, including roles of the cytoskeleton, lipids, adhesion molecules, and intrinsic endocytic protein interactions.Acknowledgments

We thank C. Qiao, V. Charles, G. Handy, L. Westhoff, A. Silveira, and M. Quiñones-Frías for technical support, and members of the Kaeser and Rodal laboratories for insightful discussions. We thank P. De Camilli for anti-Dynamin-1, anti-PIPK1γ and anti-AP-180 antibodies, D. Dickman for anti-Dynamin and anti-EndoA antibodies, N. Harden for anti-Pak antibodies, A.M.J.M. van den Maagdenberg for Ca_V_2.1 floxed mice, T. Schneider for Ca_V_2.3 floxed mice, and Troy Littleton for *brp* mutant flies. This work was supported by grants from the NIH (R01MH113349 and R01NS083898 to PSK, R01NS116375 to AAR, T32007292 to KMDLG, S10 OD034223 for the Abberior Facility Line STED microscope, and K99NS129959 to JE-M), from the NSF (Brandeis Materials Research and Engineering Center NSF-DMR 2011846), and from Harvard Medical School (Harvard/MIT Joint Research Grant in Basic Neuroscience to PSK and AAR). JE-M was supported by an Alice and Joseph E. Brooks postdoctoral fellowship from Harvard Medical School. We used fly stocks from the Bloomington *Drosophila* Stock Center (NIH P40OD018537) and antibodies from the Developmental Studies Hybridoma Bank, generated by the NICHD of the NIH. We acknowledge the Harvard Neurobiology Imaging Facility and Brandeis University Light Microscopy Core Facility Imaging Facility (RRID:SCR_025892).

## Author contributions

Conceptualization: JE-M, SJD, PSK and AAR; Methodology: JE-M, SJD; Investigation: JE-M, SJD, and KMDLG; Formal Analysis: JE-M, SJD and KMDLG; Writing-Original Draft: JE-M, SJD, PSK, AAR; Writing - Review and Editing: JE-M, SJD, KMDLG, PSK, AAR; Supervision: PSK, AAR; Funding Acquisition: JE-M, PSK, AAR. Experiments in mouse hippocampal neurons were conducted at Harvard Medical School and experiments in *Drosophila* were conducted at Brandeis University.

## Materials availability

Mouse lines, *Drosophila* lines, plasmids and antibodies will be shared upon request within the limits of the existing material transfer agreements and as long as they are available. Requests for resources and reagents should be directed to the corresponding authors.

## Data availability

Data points generated for this study are included in the figures. Additional data are available from the corresponding authors upon request.

## MATERIALS AND METHODS

### Key resource table

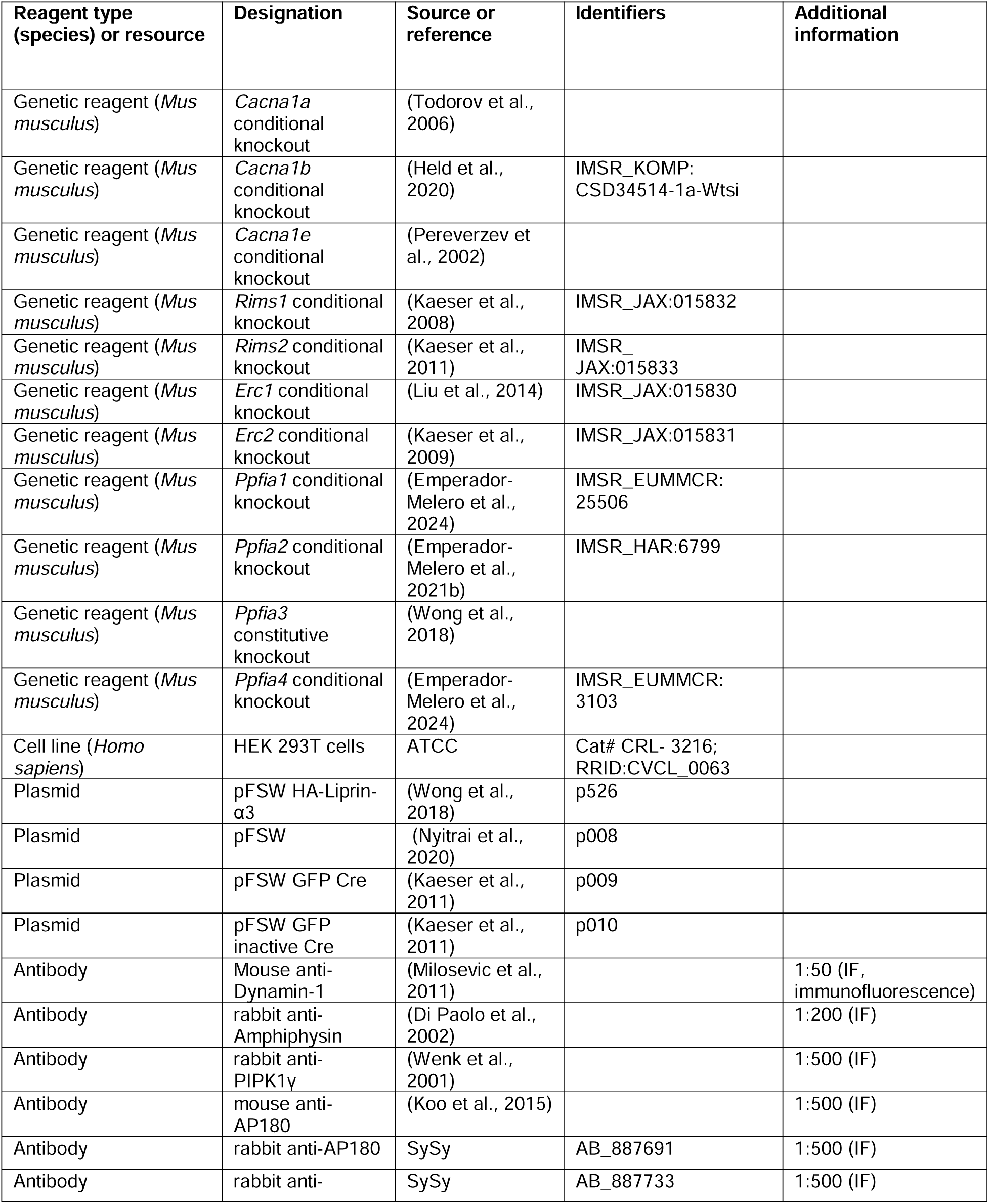
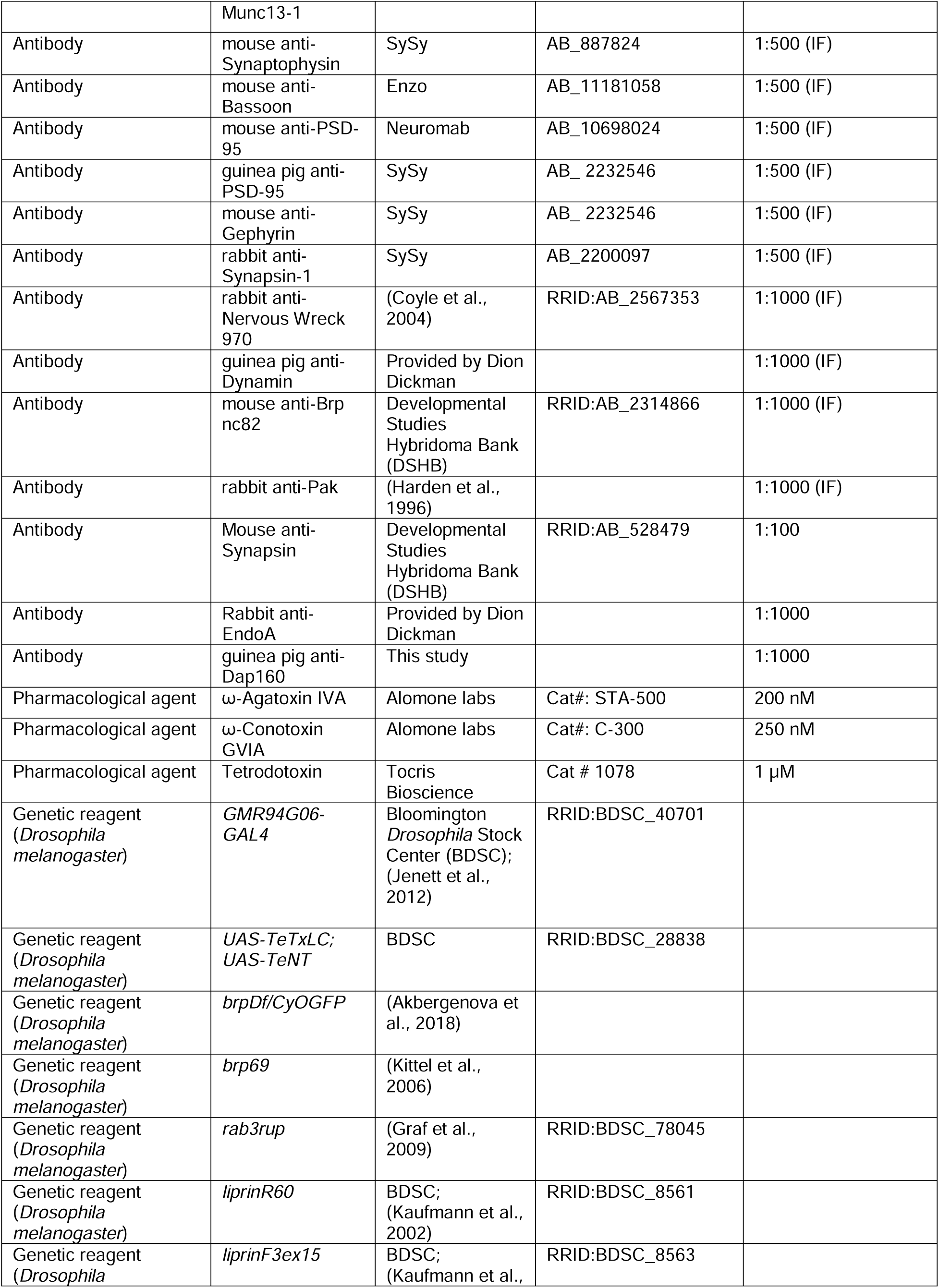
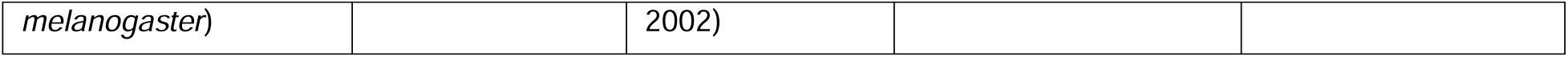

### Mouse lines

The triple mutant mice for ablating Ca_V_2.1 (targeting *Cacna1a* (Todorov et al., 2006)), Ca_V_2.2 (targeting *Cacana1b* (Held et al., 2020); RRID: IMSR_KOMP: CSD34514-1a-Wtsi) and Ca_V_2.3 (targeting *Cacana1e* (Pereverzev et al., 2002)) have been previously described (Chin and Kaeser, 2024; Held et al., 2020). Quadruple mutant mice for ablating RIM1 (targeting *Rims1* to remove RIM1α and RIM1β (Kaeser et al., 2008); RRID: IMSR_JAX:015832), RIM2 (targeting *Rims2* to remove RIM2α, RIM2β and RIM2γ (Kaeser et al., 2011); RRID: IMSR_JAX:015833), ELKS1α (targeting *Erc1* to remove ELKS1αA and ELKS1αB (Liu et al., 2014); RRID: IMSR_JAX:015830) and ELKS2α (targeting *Erc2* to remove ELKS2αA and ELKS2αB (Kaeser et al., 2009); RRID: IMSR_JAX:015831) were previously described (Emperador-Melero et al., 2024; Tan et al., 2022; Wang et al., 2016). Quadruple mutant mice for ablating Liprin-α1 (targeting *Ppfia1* (Emperador-Melero et al., 2024); RRID: IMSR_EUMMCR:25506), Liprin-α2 (targeting Ppfia2 (Emperador-Melero et al., 2021b); RRID: IMSR_HAR:6799), Liprin-α3 (targeting *Ppfia3* (Wong et al., 2018)) and Liprin-α4 (targeting *Ppfia4* (Emperador-Melero et al., 2024); RRID: IMSR_EUMMCR:3103) were previously described (Emperador-Melero et al., 2024). Mouse lines were maintained as homozygotes for the respective mutations, and offsprings were weaned at 21 to 28 days. Mice were kept separated by sex or housed as breeding pairs in a room with a regular dark light cycle and set to 22 °C (range 20 °C to 24 °C) and 50% humidity (range 35% to 70%). Experiments were approved by the Harvard University Animal Care and Use Committee.

### Primary mouse hippocampal cultures

Primary hippocampal cultures were prepared using previously established protocols (Emperador-Melero et al., 2024; Held et al., 2020; Nyitrai et al., 2020; Wang et al., 2016; Wong et al., 2018). Hippocampi dissected from newborn pups (P0-P1) were digested in papain and dissociated. Dissociated cells were plated onto 12 mm, #1.5 glass coverslips in “plating medium” that contained Minimum Essential Medium (MEM) supplemented with 0.5% glucose, 0.02% NaHCO3, 0.1 mg/mL transferrin, 10% Fetal Select bovine serum (Atlas Biologicals FS-0500-AD), 2 mM L-glutamine, and 25 mg/mL insulin. Twenty-four hours after plating, the medium was exchanged with “growth medium” that contained MEM with 0.5% glucose, 0.02% NaHCO3, 0.1 mg/mL transferrin, 5% Fetal Select bovine serum (Atlas Biologicals FS-0500-AD), 2% B-27 supplement, and 0.5 mM L-glutamine. At 48 to 60 hours after plating, cytosine β-D-Arabinofuranoside (AraC) was added at a final concentration of 2-6 µM depending on glial growth. Cells were kept in an incubator at 37 °C until DIV15 to DIV16. For activity blockade, a cocktail of drugs containing Tetrodotoxin (TTX), ω-Agatoxin IVA (ω-Aga) and ω-Conotoxin GVIA (ω-Cono) was added at DIV3 to a final concentration of 1 µM, 250 nM and 200 nM, respectively, and supplemented every three days. KCl was added to a final concentration of 50 mM from a 10X stock for 30 seconds before fixation.

### Cell lines

HEK293T cells, an immortalized cell line of female origin, were purchased from ATCC (CRL-3216, RRID: CVCL_0063) as a mycoplasma-free cell line. These cells were maintained using established methods (Emperador-Melero et al., 2024; Held et al., 2020; Nyitrai et al., 2020; Wang et al., 2016; Wong et al., 2018). They were expanded and stored in liquid nitrogen until use. After thawing, cells were grown in Dulbecco’s Modified Eagle Medium (DMEM) with 10% Fetal bovine serum (Atlas Biologicals F-0500-D) and 1% Penicillin-Streptomycin. HEK293T cells were passaged at ratios between 1:3 and 1:10 up to ∼25 times, when they were replaced with a freshly thawed batch of cells.

### Production of lentiviruses and transduction of primary mouse neurons

Lentiviruses were prepared using established methods (Emperador-Melero et al., 2024; Held et al., 2020; Nyitrai et al., 2020; Wang et al., 2016; Wong et al., 2018). HEK293T cells were transfected using the Ca^2+^ phosphate method with a combination of three packaging plasmids (REV, RRE and VSV-G), and a lentiviral plasmid encoding either GFP-tagged Cre recombinase (Kaeser lab plasmid code p009), an inactive version of Cre (p010), Liprin-α3 (p526) or a lentivirus without an insert in the multiple cloning site (p008) at a molar ratio 1:1:1:1 and with a total amount of ∼4 μg DNA per T25 flask. At 20 to 30 hours after transfection, the medium was changed to “growth medium” and 48 to 60 hours after transfection the supernatant was collected. Viral transductions were done either with freshly collected viral supernatant or with snap-frozen supernatant stored at -80 °C. Neuronal cultures were transduced with lentiviruses expressing Cre or inactive Cre at DIV2 for Ca_V_2 ablation (Held et al., 2020), DIV5 for RIM+ELKS ablation (Emperador-Melero et al., 2024; Tan et al., 2022; Wang et al., 2016), or DIV7 for Liprin-α ablation (Emperador-Melero et al., 2024) following protocols established before. Transduction efficiency was monitored via the presence of nuclear GFP fluorescence, and only cultures in which no neurons without nuclear green fluorescence were readily detected were used for experiments. Liprin-α mutant neuronal cultures were additionally transduced at DIV1 with either lentivirus expressing either HA-tagged Liprin-α3 (p526; for control^L1-L4^ neurons) or lentivirus without an insert in the multiple cloning site (p008; for cQKO^L1-L4^ neurons). pFSW HA-Liprin-α3 (p526) corresponds in sequence and numbering to NCBI Reference Sequence: NP_001257914.1 with the addition of an N-terminal HA-tag and a short linker (sequence: M-YPYDVPDYA-GAPS-C_3_…C_1192_) and has been described before (Emperador-Melero et al., 2024, 2021b; Wong et al., 2018).

### Immunofluorescence staining for STED and confocal microscopy of cultured hippocampal neurons

Neurons were antibody-stained using previously established protocols (Emperador-Melero et al., 2024, 2021b; Held et al., 2020; Wong et al., 2018). They were fixed at DIV15 to DIV16 in PBS containing 2% PFA for 10 minutes at room temperature. For cultures stimulated with KCl, 50 mM KCl was also included in the fixation solution after the 30 s depolarization in described below. Neurons were blocked and permeabilized in blocking solution (PBS, 3% BSA, 0.1% Triton X-100) for 1 hour at room temperature and incubated with primary and secondary antibodies, each overnight at 4 °C, in blocking solution. Three 5-minute washes with PBS were performed after each antibody incubation. The cells were post-fixed in 4% PFA for 10 min, washed in PBS, and mounted using mounting medium. Primary antibodies used were: mouse anti-Dynamin-1 (Kaeser lab antibody code A242, 1:50, obtained from P. De Camilli, knockout-validated for immunostaining in (Milosevic et al., 2011)), rabbit anti-Amphiphysin (A244, 1:200, obtained from P. De Camilli, knockout-validated for immunostaining in (Di Paolo et al., 2002); rabbit anti-PIPK1γ (A168, 1:500, obtained from P. De Camilli (Wenk et al., 2001)); mouse anti-AP-180 (A246, 1:500, obtained from P. De Camilli, knockout-validated for immunostaining in (Koo et al., 2015); rabbit anti-AP-180 (A219, 1:500; RRID: AB_887691); rabbit anti-Munc13-1 (A72, 1:500; RRID: AB_887733); mouse anti-Synaptophysin (A100, 1:500; RRID: AB_887824); mouse anti-PSD-95 (A152, 1:500; RRID: AB_10698024); mouse anti-Bassoon (A85, 1:500; RRID: AB_11181058); guinea pig anti-PSD-95 (A5, 1:500, RRID: AB_2619800); rabbit anti-Synapsin-1 (A30, 1:500, RRID: AB_2200097); mouse anti-Gephyrin (A8; 1:500, RRID: AB_2232546). Secondary antibodies used were: goat anti-guinea pig Alexa Fluor 633 (S34; 1:250, RRID: AB_2535757), goat anti-mouse IgG2a Alexa Fluor 555 (S20; 1:250, RRID: AB_2535776), goat anti-rabbit Alexa Fluor 488 (S5; 1:250, RRID: AB_2576217), goat anti-rabbit Alexa Fluor 555 (S22; 1:250, RRID: AB_2535849), goat anti-mouse Alexa Fluor 488 (S4; 1:250, RRID: AB_2534088), goat anti-rabbit Alexa Fluor 633 (S33; 1:250, RRID: AB_2535731) and goat anti-guinea pig Alexa Fluor 555 (S23; 1:250, RRID: AB_2535856).

### Image acquisition and analyses for confocal and STED microscopy of cultured hippocampal neurons

Images were acquired as previously established (Chin and Kaeser, 2024; Emperador-Melero et al., 2024, 2021b, 2021a). A Leica SP8 Confocal/STED 3X microscope equipped with an oil-immersion 100x objective (1.44 NA), a white laser, STED gated detectors, and 592, 660 and 770 nm depletion lasers was used to acquire 2048 pixels x 2048 pixels large images (pixel size of 22.7 nm x 22.7 nm) that contained hundreds of synapses. Triple confocal scans for Synaptophysin or Synapsin, an active zone (Munc13-1 or Bassoon) or postsynaptic density (PSD-95) marker and a protein of interest were followed by double-color STED scans for the active zone or postsynaptic density marker and the protein of interest. The exceptions were (1) combinations containing Amphiphysin and PIPK1γ in Fig. 1, where Synaptophysin was also imaged in STED, and (2) combinations containing Amphiphysin, PIPK1γ or AP-180 in Fig. 6, where an antibody against Gephyrin was also added and imaged in confocal and STED modes as well. Acquisition settings for a given staining and channel were identical for all images within a batch of culture in which all conditions from one experiment were compared. To quantify STED images, side-view synapses were selected by an experimenter blind to the protein of interest. Side-view synapses were defined as those containing a vesicle cloud of 250 nm or more in width from the marker to the inside of the presynaptic terminal and with a bar-like Munc13-1, Bassoon or PSD-95 structure along its edge. A 750-nm long, rectangular area of interest with a width exceeding that of the active zone or postsynaptic marker by up to five pixels on each side was drawn perpendicular to the marker and across its center. After applying a 5-pixel rolled average to the protein of interest and marker, line profiles of individual synapses were aligned to the peak of the active zone or postsynaptic marker and averaged. Next, the position of the maximum value of the endocytic protein relative to the maximum value of the marker was calculated and plotted for each synapse. The maximum fluorescence value of the endocytic protein within 136 nm relative to the peak fluorescence of the marker was also measured and plotted. For Figs. 1 and 3, this region spanned 68 nm at each side of the peak of the active zone marker. For Figs. 4 and 6, it spanned 136 nm from the PSD-95 peak towards the presynaptic bouton. These distances were chosen to match the periactive zone area based on the position of synaptic proteins at STED resolution (Wong et al., 2018). In each culture, line profiles and peak intensities were normalized to the average signal in the condition used for comparison (defined in the corresponding figure legend). In line profile plots, peaks are below 100% because peak intensities for proteins of interest are not always at the same position. En-face synapses were selected as synapses that did not have a bar-like appearance of the active zone or postsynaptic marker; instead, the area of the marker was surrounded by Synaptophysin or Synapsin staining. The experimenter was blind to the protein of interest during en-face synapse selection. For any given en-face synapse, only signals of the protein of interest that fell within 136 nm of the edges of the synaptic vesicle cloud were included. This area was defined by creating a binary mask for Synaptophysin or Synapsin and expanding it by 6 pixels (each 22.7 nm). Individual channels containing the marker or the protein of interest were next thresholded. For any given experiment, thresholds were defined by an experimenter blind to the condition through visual inspection of approximately ten images, and the same thresholds were then applied to all images within an experiment. After thresholding, the “analyze particles” function of Fiji was applied to detect individual objects and to measure their size, position and intensity (the intensity was measured in the original, non-thresholded image). For quantification of confocal signals, individual channels were analyzed with an automatic detection algorithm (available at https://github.com/kaeserlab/3DSIM_Analysis_CL). With this algorithm, Otsu thresholding was used to generate a presynaptic mask based on the Synaptophysin or Synapsin signals. The synaptic mask was subsequently used to quantify the fluorescence intensity levels of the protein of interest. To avoid detection artifacts, areas with somata and out-of-focus areas were not included in the areas of interest. Confocal data were normalized to the average in the condition that was used for comparison (noted in each figure legend) per culture. For representative images, a smooth filter was added in some cases, brightness and contrast were linearly adjusted, and images were interpolated. Identical adjustments were applied to representative images of the same channel within an experiment. Quantifications were performed on original images without brightness and contrast adjustments and without background subtraction or resampling. Data were acquired and analyzed by an experimenter blind to genotype.

### Drosophila strains

Flies were cultured using standard media and techniques. All flies were raised at 25 °C. Fly strains used in this work were: GMR94G06-GAL4 RRID:BDSC_40701, UAS-TeTxLC (aka UAS-TeNT, RRID:BDSC_28838), *brp*^Df^ (Akbergenova 2018), *brp*^69^ (Kittel 2006), *rab3^rup^* (RRID:BDSC_78045 (Graf 2007)), *liprin^R60^* (RRID:BDSC_8561, (Kaufmann, 2002)), *liprin*^F3ex15^ (RRID:BDSC_8563 (Kaufmann, 2002)).

### Generation of polyclonal antibodies

*Drosophila* antigens for producing anti-Dap160, anti-Dynamin, and anti-EndoA antisera were produced in lab and sent to Cocalico Biologicals, Inc. (Denver, PA, USA) for injection into two guinea pigs each, and antisera were harvested. Specificity of the sera was assessed by staining NMJs of control larvae and of larvae with RNAi against the protein of interest expressed pan-neuronally by C155-Gal4 (Fig. 2 - figure supplement 1).

Guinea pig anti-Dap160 polyclonal antisera were raised against a recombinant protein fragment of Drosophila Dap160 SH3 domains with a 6x-His N-terminal tag (sequence: MGGSHHHHHH-GMASMTGGQQMGRDLYDDDDKDRWGSTGSSSAWEETGTTVTDPYAVASNDISALAAPAVDLGGPAPEGFVKYQAVYEFNARNAEEITFVPGDIILVPLEQNAEPGWLAGEINGHTGWFPESYVEKLEVGEVAPVAAVEAPVDAQVATVADTYNDNINTSSIPAASADLTAAGDVEYYIAAYPYESAEEGDLSFSAGEMVMVIKKEGEWWTGTIGSRTGMFPSNYVQKADVGTASTAAAEPVESLDQETTLNGNAAYTAAPVEAQEQVYQPLPVQEPSEQPISSPGVGAEEAHEDLDTEVSQINTQSKTQSSEPAESYSRPMSRTSSMTPGMRAKRSEIAQVIAPYEATSTEQLSLTRGQLIMIRKKTDSGWWEGELQAKGRRRQIGWFPATYVKVLQGGRNSGRNTPVSGSRIEMTEQILDKVIALYPYKAQNDDELSFDKDDIISVLGRDEPEWWRGELNGLSGLFPSNYVGPFVTSGKPAKANGTTKK). The protein was expressed in bacteria and purified with a Ni^2+^-column followed by gel filtration. Sera were not pooled, and experiments reported here used the pre-production test bleed.

Guinea pig anti-Dynamin antisera were provided by D. Dickman; they were raised against a recombinant peptide conjugated to KLH (sequence: (C)-RPGGSLPPPMLPSRR).

Rabbit anti-EndoA polyclonal antisera were provided by D. Dickman; they were raised against a recombinant protein with a 6x-His N-terminal tag expressed in bacteria and purified with a Ni^2+^-column (sequence: MHHHHHH-KEFLQPNPTARAKMAAVKGISKLSGQAKSNTYPQPEGLLAECMLTYGKKLGEDNSVFAQALVEFGEALKQMADVKYSLDDNIKQNFLEPLHHMQTKDLKEVMHHRKKLQGRRLDFDCKRRRQAKDDEIRGAEDKFGESLQLAQVGMFNLLENDTEHVSQLVTFAEALYDFHSQCADVLRGLQETLQEKRSEAESRPRNEFVPKTLLDLNLDGGGGGLNEDGTPSHISSSASPLPSPMRSPAKSMAVTPQRQQQPCCQALYDFE).

### Immunostaining of Drosophila NMJs

For analyses of NMJ morphology and protein localization, flies were maintained at low density at 25 °C. Wandering third instar larvae were dissected in Ca^2+^-free HL3.1 saline (70 mM NaCl, 5 mM KCl, 10 mM MgCl2, 10 mM NaHCO3, 5 mM trehalose, 5 mM HEPES, 115 mM sucrose (Feng et al., 2004)) and fixed for 20 min in HL3.1 containing 4% PFA. Fixed larvae were incubated in “blocking solution” containing 3% BSA, 0.1% Triton X-100 in PBS for 30 to 60 min and incubated with primary antibody in blocking solution either overnight at 4 °C or for 2 hours at room temperature. Samples were rinsed three times in a “washing buffer” containing 0.1% Triton in PBS followed by three ten-minute washes in washing buffer. Samples were incubated for one hour at room temperature with dye-conjugated secondary antibodies diluted to 1:250 in washing buffer, followed by washing as after primary antibody incubation. Larvae were mounted in Abberior Mount Liquid mounting medium. Primary antibodies used were rabbit anti-Nwk 970 (RRID:AB_2567353, knockout validated for immunostaining in (Coyle et al., 2004)), guinea pig anti-Dynamin (gift from D. Dickman, validated for immunostaining by RNAi, Figure 2 - figure supplement 1), guinea pig anti-Dap160 (validated for immunostaining by RNAi, Figure 2 - figure supplement 1), rabbit anti-endophilin (gift from D. Dickman, validated for immunostaining by RNAi, Figure 2 - figure supplement 1), mouse anti-Dynamin (RRID:AB_397640, BD Biosciences Clone 41, validated for immunostaining by RNAi (Kasprowicz et al., 2014)), mouse anti-BRP (RRID:AB_2314866, DSHB clone nc82, validated for immunostaining by RNAi (Wagh et al., 2006)), rabbit anti-Pak (gift of N. Harden, mutant-validated for immunostaining (Harden et al., 1996)). Triple-labeling of Pak, FasII, and Brp was performed as follows: Samples were incubated with anti-Pak and anti-FasII primary antibodies overnight at 4 °C, followed by washing, species specific secondary antibody incubation, and further washing using the same buffers and procedures as described above. Next, Brp was labeled using anti-Brp (nc82; conjugated directly to Alexafluor-568, using a commercial antibody labeling kit (Thermofisher) for four hours at room temperature, followed by three ten-minute washes.

### NMJ image acquisition and processing

Confocal images of NMJs were acquired at room temperature with a Zeiss 880FAS microscope in SR mode, using a 63X (NA1.4) oil immersion objective and Zen Black software. All raw image stacks were processed in Zen Blue to construct Airyscan images using 3D Airyscan processing with automatic settings. Lateral and axial resolution were estimated to be ∼175 nm and ∼400 nm, respectively, by imaging Tetraspeck beads (Invitrogen) at 560 nm. To prepare images for analyses, we excluded regions that were obscured by axon bundles or contained Type 1s terminals.

To measure intensities in whole boutons, a 3D presynaptic mask was generated as follows. We produced a normalized sum image of Nwk, Dynamin, and Brp signals (except Fig. 6, which lacks Brp staining) by dividing each channel by its mean and summing the channels. Sum images were then gaussian filtered (sigma = 5 pixels for all experiments except Fig. 9C-G, where sigma = 4) and thresholded by intensity by a previously established algorithm (Li and Tam, 1998) (except Fig. 9C-G, which used Otsu thresholding (Otsu, 1979)). The binary mask was eroded by 4 pixels (1 pixel for Fig. 9C-G). Mask settings were manually confirmed to accurately represent the 3D volume across images, and identical settings were used within an experiment. For all images, the background was subtracted using the rolling ball method with a radius of 50 pixels, and signal intensities were measured in 3D using a custom FIJI script (available at https://github.com/rodallab/nmj-measurement). For measurements of protein levels and polarization at the periactive zone, we used custom FIJI and Python scripts described in detail (Del Signore et al., 2023) and available at https://github.com/rodallab/paz-analysis. First, maximum intensity projections were made of the upper half of NMJ terminals, to analyze a single plasma membrane surface. Second, a 2D mask of the total presynaptic area was generated by summing all channels in each image and then thresholded by intensity using a previously established algorithm (Li and Tam, 1998). Third, a periactive zone mesh composite image was created by subtracting the Brp signal from the sum of Nwk and Dyn signals, with the following exceptions: For Fig. 6, only the Nwk was used. For Fig. 9L-N, Brp and PAK signals were subtracted from FasII to create the composite mask. Fourth, periactive zone units were detected as local intensity minima and then expanded by the seeded region growing algorithm (via the IJ-Plugins Toolkit). The thresholds for minima detection were computed automatically and periactive zones detected at the edge of boutons (defined as having a mean Euclidean distance map score of less than 7.5 pixels) were excluded from analysis as these are not planar (Del Signore et al., 2023). As above, settings were applied identically within an experiment. Image analyses settings were computed without manual user intervention and data acquisition and analyses were not blinded. For representative images, brightness and contrast were linearly adjusted and applied identically to representative images of the same channel within an experiment.

### Electrical stimulation of Drosophila NMJs

Control *white*^1118^ animals were dissected at room temperature in HL3.1 with 0 mM extracellular Ca^2+^, and motor axons were cut close to the ventral ganglion. Prior to electrical stimulation, the tissue sample was washed three times with 1 mL of HL3.1 solution containing 2 mM Ca^2+^ and 7 mM glutamate (70 mM NaCl, 5 mM KCl, 10 mM MgCl_2_, 10 mM NaHCO_3_, 5 mM trehalose, 5 mM HEPES, 115 mM sucrose, 2 mM CaCl_2_, 7 mM monosodium glutamate). The axon bundle innervating segment A3 was sucked into a pipette and 40 Hz stimulation was administered for 3 minutes (5 V for 0.5 ms per stimulus) at room temperature. The stimulus was administered through an A-M system 2100 stimulator using ADinstruments lab chart software. Stimulation was monitored by observing muscle contraction of the desired abdominal segment through the eyepiece. For stimulation experiments, muscles 6/7 and muscle 4 were imaged and analyzed together. As controls, comparisons were made to NMJs from the contralateral, unstimulated muscles. After stimulation, the preparation was stretched and fixed within 30 seconds with 4% PFA for 15 minutes, and samples were processed for immunohistochemistry as described above.

### Statistics

Data are shown as mean ± SEM. For analyses of STED images of hippocampal synapses, the sample size is the number of analyzed synapses. For analyses of confocal images of hippocampal synapses, the sample size is the number of analyzed images. For analyses of *Drosophila* NMJs, the sample size is the number of analyzed NMJ terminals. Sample sizes and statistical tests are included in each figure legend. Significance is reported as * p < 0.05, ** p < 0.01, or *** p < 0.001 and was assessed using parametric (t-test or one-way ANOVA) or non-parametric (Mann-Whitney U or Kruskal-Wallis) tests depending on whether assumptions of normality and homogeneity of variances were met (assessed using Shapiro or Levene’s tests, respectively). Tukey-Kramer or Holm corrections for multiple testing were applied. In Fig. 1 and Fig. 1 - figure supplement 1, post-hoc comparisons between all groups were performed, and only significance relative to the untreated condition is reported. For STED images, statistical analyses were performed on the peak values of the line profiles. Statistical analyses were performed in R.

---

**Figure 1 - figure supplement 1.**
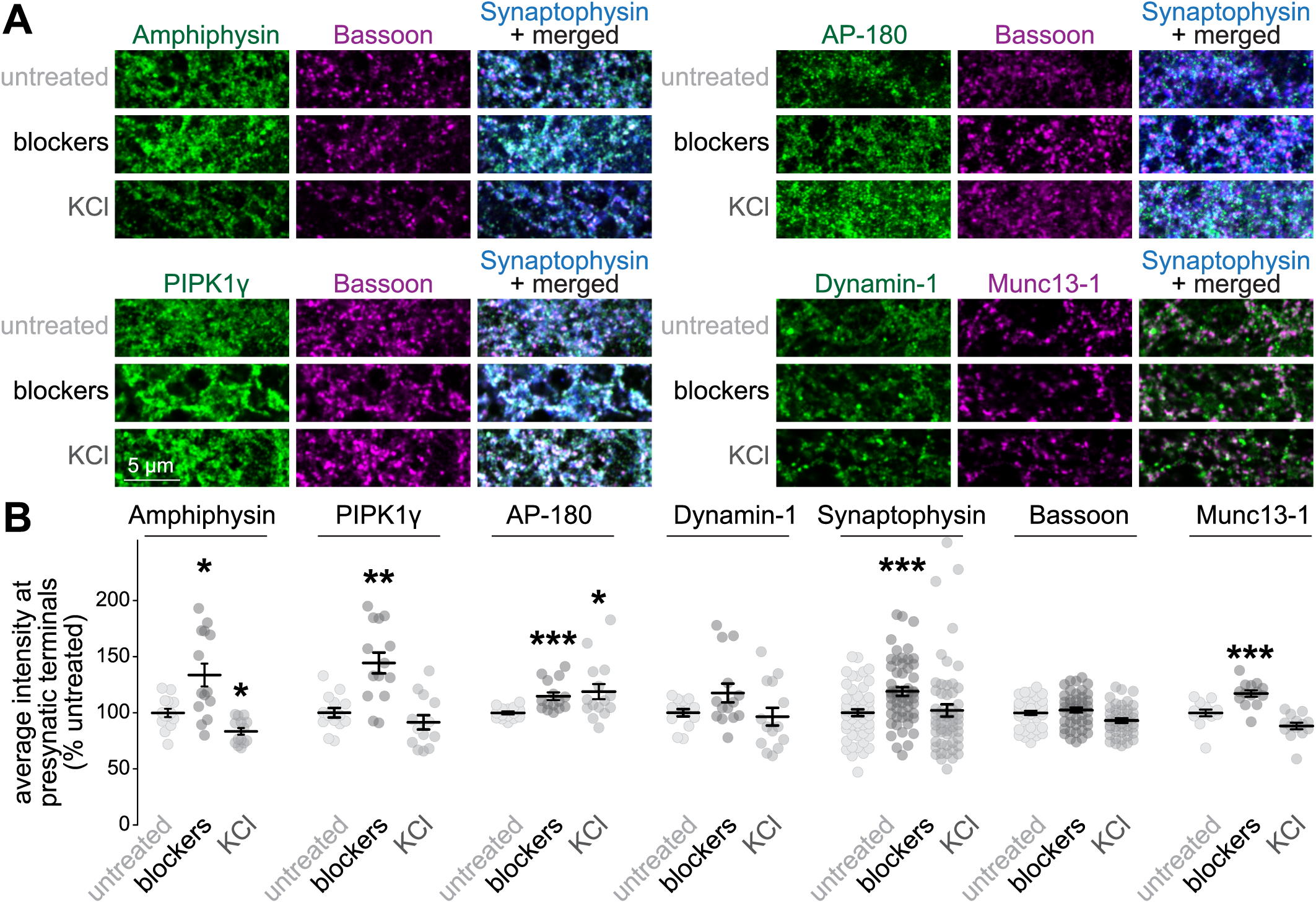
Confocal microscopic analyses of synapses after chronic silencing or acute depolarization of mouse hippocampal neurons. (**A, B**) Example confocal images (A) and quantification of the average intensities (B) of Amphiphysin, PIPK1γ, AP-180, Dynamin-1, Bassoon and Munc13-1 at synapses identified as Synaptophysin puncta. Intensities are normalized to the average signals in the untreated conditions per culture; n in B (images/cultures): Amphiphysin, 14/3; PIPK1γ, untreated 14/3, blockers 14/3, KCl 13/3; AP-180, 15/3; Dynamin-1, 14/3; Synaptophysin, untreated 57/3, blockers 57/3, KCl 56/3; Bassoon, untreated 43/3, blockers 43/3, KCl 42/3; Munc13-1, 14/3. The increase in Munc13-1 upon chronic silencing, which we previously reported (Held et al., 2020), and of Synaptophysin, may reflect a homeostatic adaptation. The decrease in Amphiphysin upon KCl stimulation may reflect a redistribution of this protein during prolonged stimulation. Data are mean ± SEM; *p < 0.05, **p < 0.05, ***p < 0.001 compared to the untreated condition determined by one-way ANOVA followed by a Tukey-Kramer post hoc tests for Bassoon or Kruskal-Wallis followed by Holm post hoc tests for Amphiphysin, PIPK1γ, AP-180, Dynamin-1, Synaptophysin and Munc13-1.

**Figure 1 - figure supplement 2.**
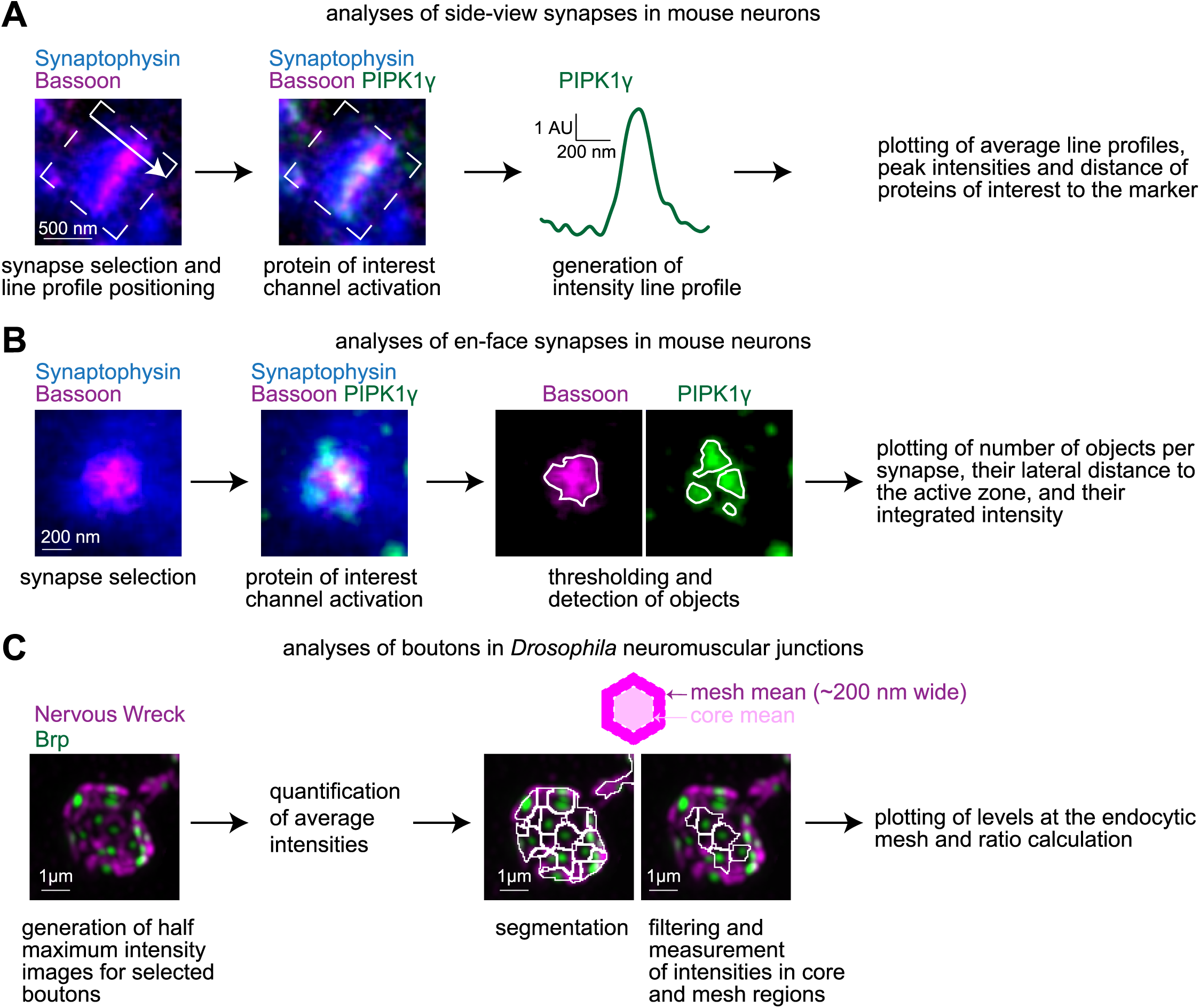
Workflows for STED analyses in mouse hippocampal neurons and for confocal analyses at *Drosophila* neuromuscular junctions. (**A**) Workflow for the analyses of side-view synapses of mouse hippocampal neurons, showing an example synapse immunostained for the active zone marker Bassoon (imaged in STED), PIPK1γ (imaged in STED) and Synaptophysin (imaged in confocal). Synapse selection and placement of an area of interest (white rectangle with line profile direction indicated by arrow) perpendicular to the marker (Bassoon, in this example) is done by an experimenter blind for the protein of interest (PIPK1γ, in this example). Next, the protein of interest channel is activated, and the profile is generated. Finally, the average line profiles, peak intensities and distance of proteins of interest to the marker are plotted. (**B**) Workflow for the analyses of en-face synapses of mouse hippocampal neurons, showing an example synapse immunostained for Bassoon (imaged in STED), PIPK1γ (imaged in STED) and Synaptophysin (imaged in confocal). First, synapse selection is done by an experimenter blind for the protein of interest. Next, the channel of the protein of interest is activated, and objects containing endocytic proteins and the marker are identified in the respective channels using an algorithm. Finally, the number of objects per synapse, their lateral distance to the active zone, and their integrated intensity are plotted. (**C**) Workflow for analyses of *Drosophila* neuromuscular junctions. Terminals are analyzed both in 3D and in 2D half-maximum intensity projections. First, the average intensities of the active zone marker (Brp in this example) and the endocytic protein (Nervous Wreck in this example) are quantified in the full 3D volume of the terminal. Next, the periactive zone levels and degree of polarization are analyzed in 2D half-maximum intensity projections. The polarization of each protein is quantified as the ratio between its average intensity at the mesh over its average intensity in the core. To conduct this analysis, segmentation into mesh and core is performed based on the difference in signal between proteins enriched in the periactive zone mesh (e.g. Nwk and Dynamin) vs. proteins enriched in the core region (e.g. Brp and Pak) as described in the methods. White lines delineate the center of the mesh regions. The mesh is ∼200 nm wide, and the core is the remaining enclosed region within the innermost bounds. The resulting ROIs are used to measure average intensities within the mesh and the core and its ratio.

**Figure 1 - figure supplement 3.**
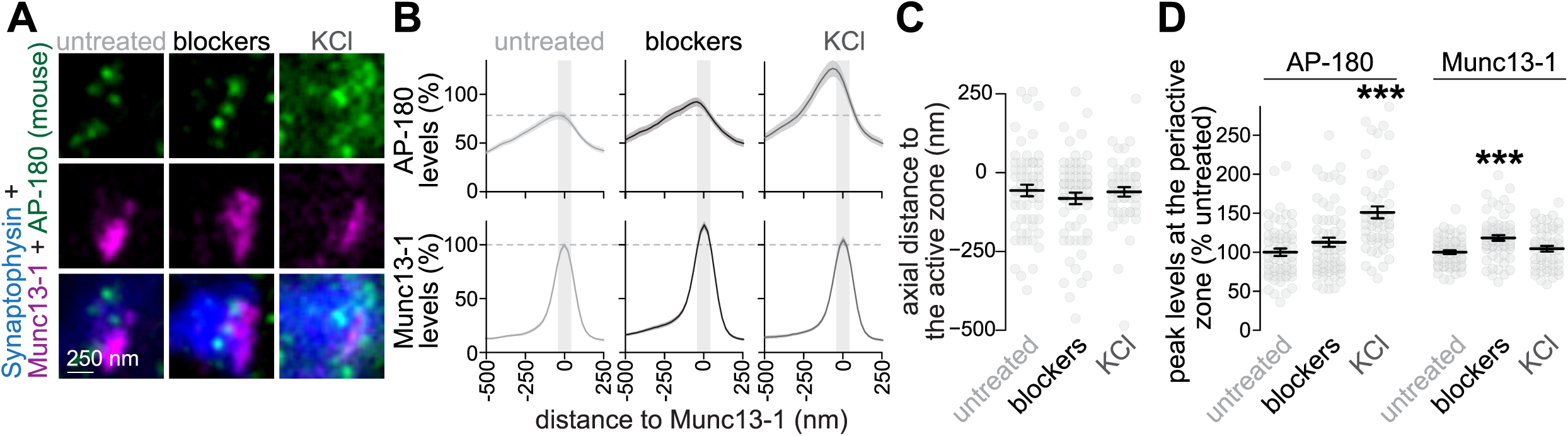
Assessment of AP-180 with alternate antibody after chronic silencing or acute depolarization of mouse hippocampal neurons. (**A, B**) Example side-view synapses (A) and average line profiles of AP-180 (antibody A246) and Munc13-1 (b). Neurons were stained for AP-180 (imaged in STED), Munc13-1 (imaged in STED), and the synaptic vesicle marker Synaptophysin (imaged in confocal). An area of interest was positioned perpendicular to the center of the Munc13-1 object, and synapses were aligned via the peak fluorescence of Munc13-1 in the average profiles. Line profiles were normalized to the average signal in the untreated condition. Dashed lines mark average levels in the untreated condition and grey shaded areas represent the active zone area; n in B (synapses/cultures): untreated, 58/3; blockers, 61/3; KCl, 50/3. **(C, D)** Quantification of the peak-to-peak distance of the active zone marker and the protein of interest (C), and of the peak levels in the periactive zone area (D). The periactive zone area is defined as an area within 68 nm on each side of the peak of the active zone marker (grey shaded areas in B); n as in B. Data are mean ± SEM; *p < 0.05, ***p < 0.001 compared to the untreated condition determined by Kruskal-Wallis followed by Holm post-hoc tests.

**Figure 1 - figure supplement 4.**
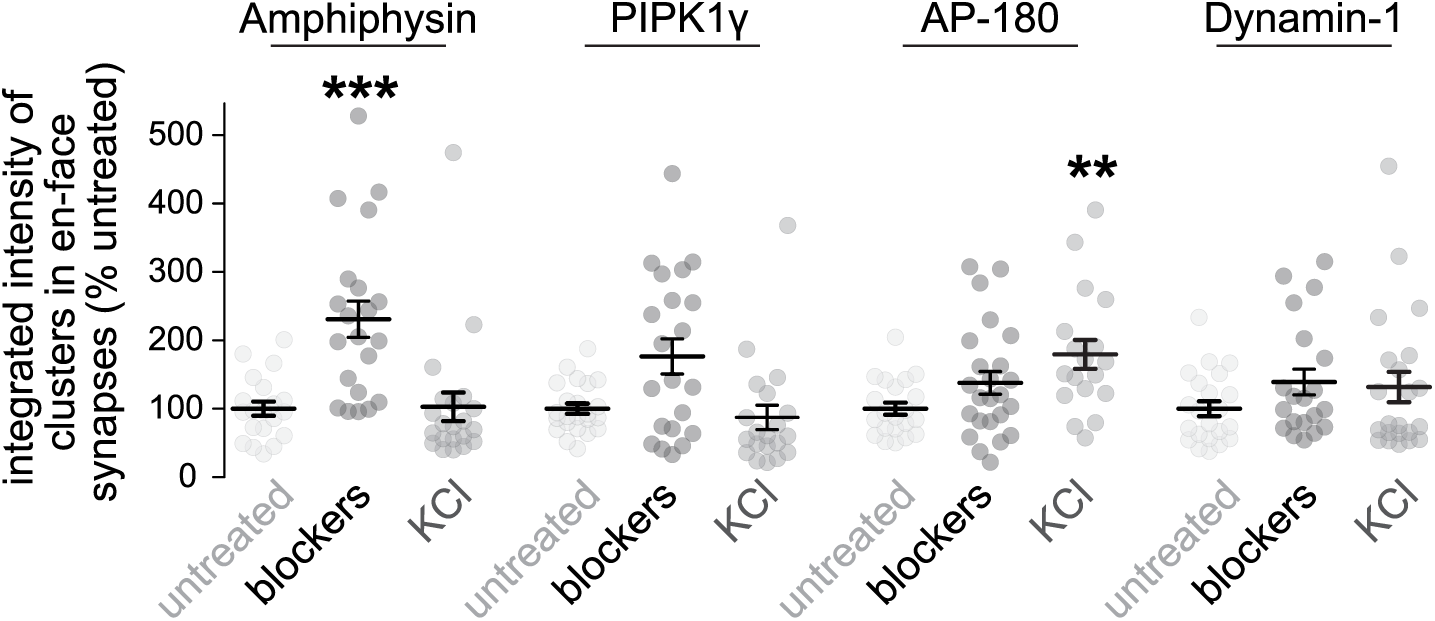
Additional analyses of en-face synapses after chronic silencing or acute depolarization of mouse hippocampal neurons. Quantification of the average integrated intensities (calculated as the object area multiplied by its average fluorescence intensity) of the Amphiphysin, PIPK1γ, AP-180 and Dynamin-1 objects detected in en-face synapses from Fig. 1L-Q. Intensities are normalized to the average signals in the untreated conditions per culture; n as in Fig. 1P. Data are mean ± SEM; **p < 0.01, ***p < 0.001 compared to the untreated condition determined by one-way ANOVA followed by a Tukey-Kramer post hoc test for AP-180 or Kruskal-Wallis followed by Holm post hoc tests for Amphiphysin, PIPK1γ and Dynamin-1.

**Figure 2 - figure supplement 1.**
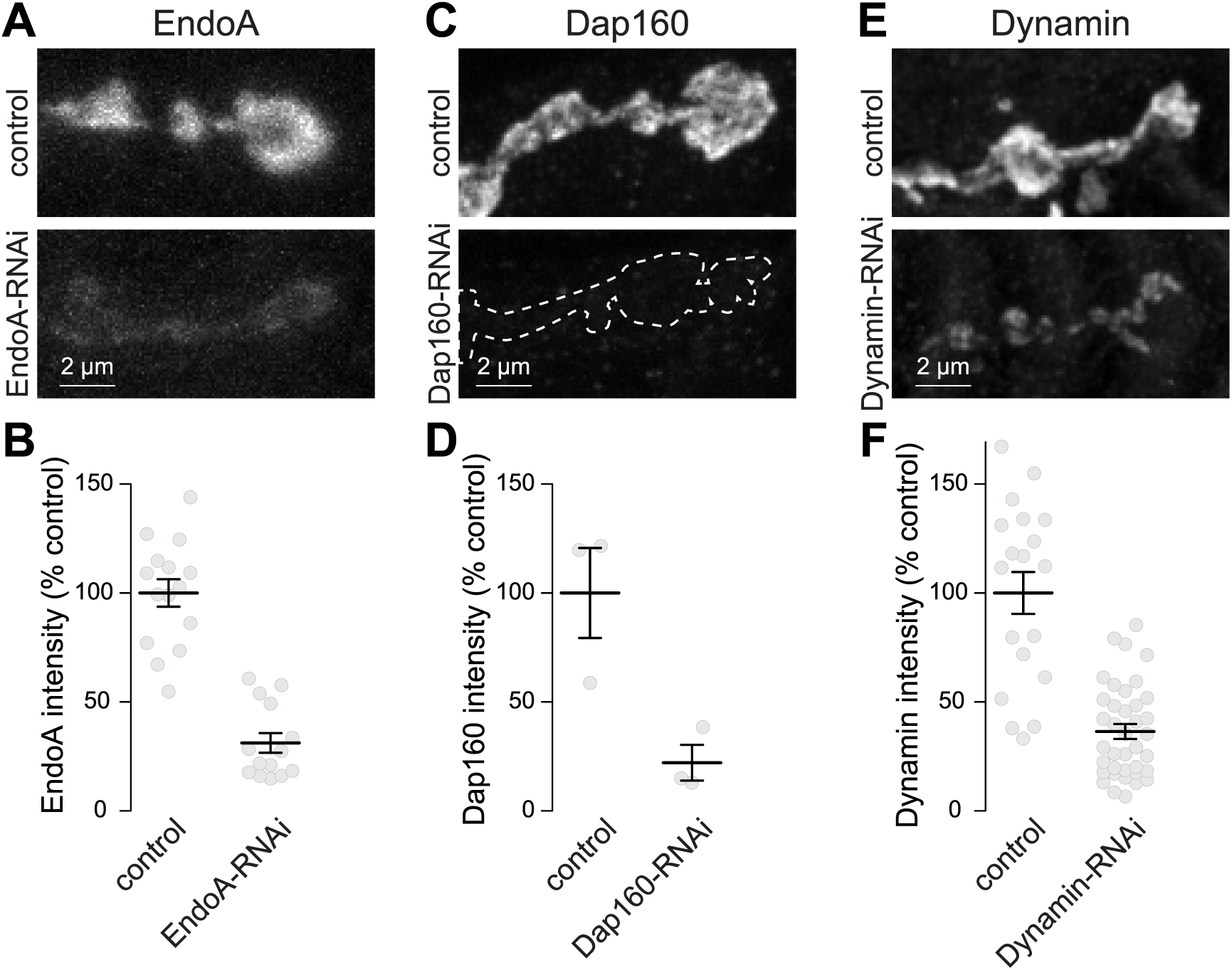
Validation of EndoA, Dap160, and Dynamin antibodies in *Drosophila* NMJs. Example confocal images and quantification of the signal of EndoA (A+B), Dap160 (C+D) or Dynamin (E+F) in NMJs expressing either driver alone (control) or an RNAi against the indicated gene. Data are expressed as the percentage of the control; n in B (NMJs/animal): control 15/3, EndoA-RNAi 14/3; D: control 3/2, Dap160-RNAi 3/2; F: control 19/6, Dyn-RNAi 38/6. Data are mean ± SEM.

**Figure 3 - figure supplement 1.**
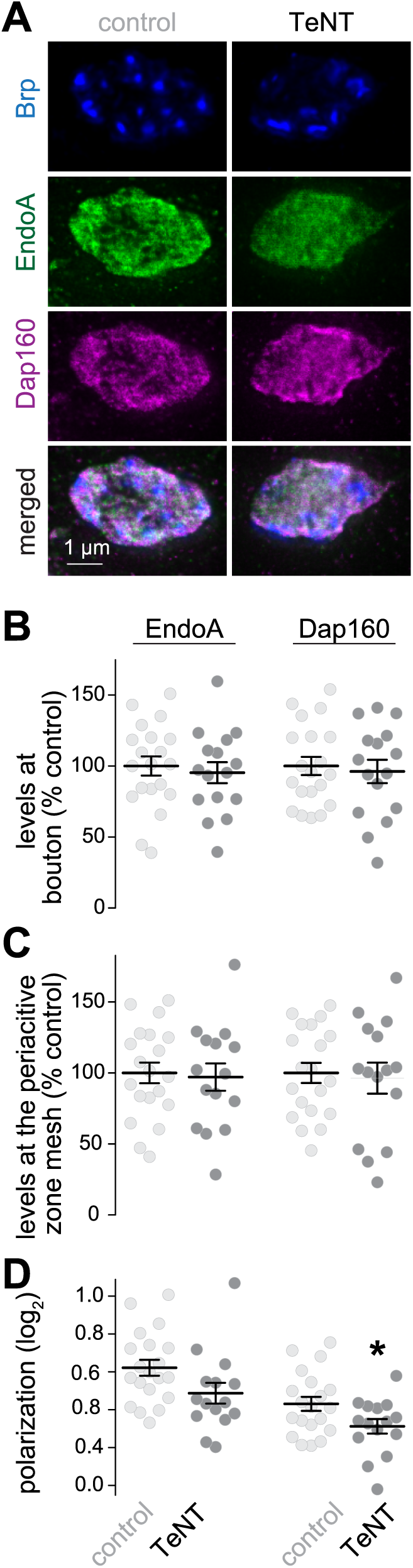
Assessment of EndoA and Dap160 after chronic silencing of *Drosophila* NMJs using STED microscopy. Example boutons with or without TeNT-expression (A), and quantification of the average fluorescence intensity of EndoA and Dap160 per bouton (B), average intensity at the periactive zone mesh (C) and the polarization within periactive zone units (D). Data in B and C are normalized to the average of the control condition; n in B (NMJ/animal): control 20/6, TeNT 16/6; C-D: control 20/6, TeNT 15/6. Data are mean ± SEM; *p < 0.05 determined by two-sided Student’s t-tests. Images acquired by STED microscopy.

**Figure 4 - figure supplement 1.**
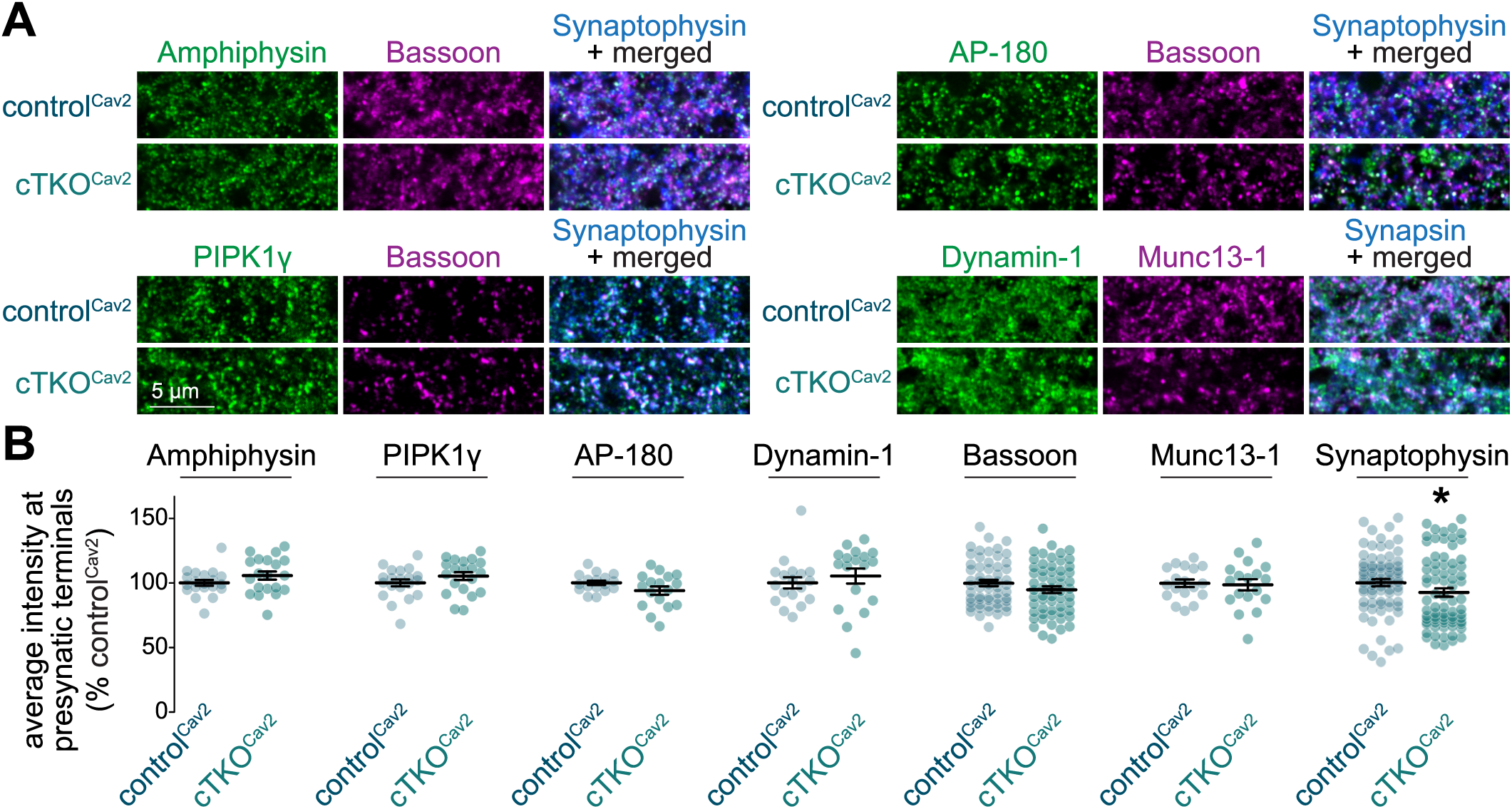
Confocal microscopic analyses of synapses after Ca_V_2 ablation in mouse hippocampal neurons. (**A, B**) Example confocal images (A) and quantification of the average intensities (B) of Amphiphysin, PIPK1γ, AP-180 and Dynamin-1 at synapses identified as Synaptophysin puncta. Intensities are normalized to the average signals in the control^Cav2^ condition per culture; n in B (images/cultures): Amphiphysin 20/3, PIPK1γ 20/3, AP-180 17/3, Dynamin-1 18/3, Bassoon 57/3, Munc13-1 18/3, Synaptophysin 75/3. Data are mean ± SEM; *p < 0.05 determined by a two-sided Student’s t-tests for Amphiphysin, PIPK1γ, Bassoon and Munc13-1, or two-sided Mann-Whitney U tests for AP-180, Dynamin-1 and Synaptophysin.

**Figure 4 - figure supplement 2.**
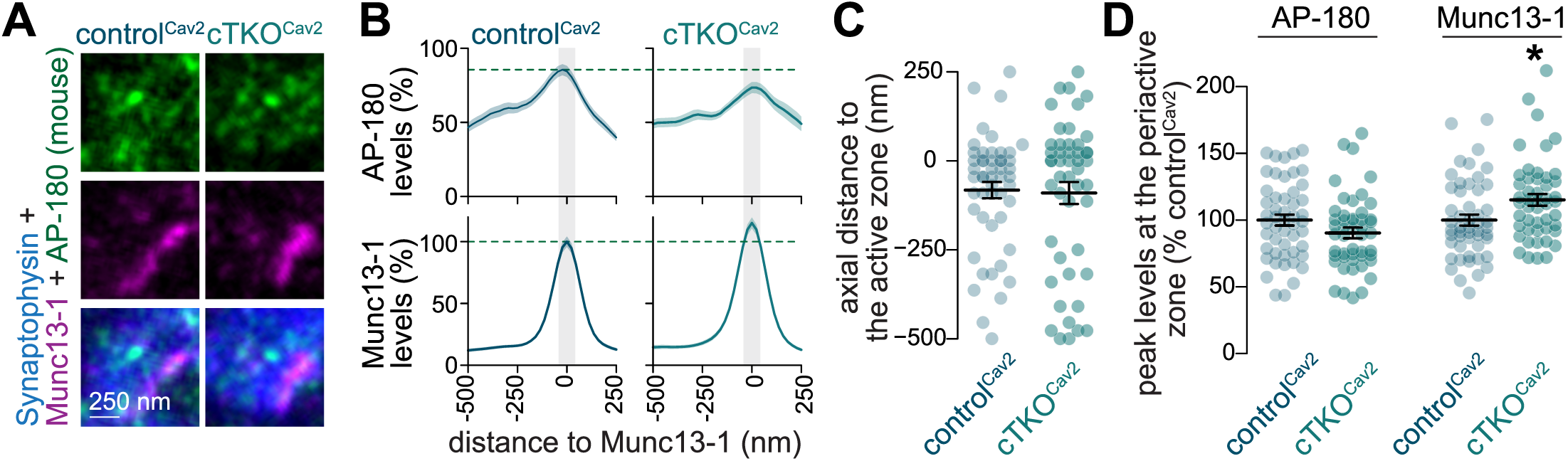
Assessment of AP-180 with an alternate antibody after Ca_V_2 ablation in mouse hippocampal neurons. (**A, B**) Example side-view synapses (A) and average line profiles (B) of AP-180 (antibody A246) and Munc13-1. Neurons were stained for AP-180 (imaged in STED), Munc13-1 (imaged in STED), and the synaptic vesicle marker Synaptophysin (imaged in confocal). An area of interest was positioned perpendicular to the center of the Munc13-1 object, and synapses were aligned via the peak fluorescence of Munc13-1 in the average profiles. Line profiles were normalized to the average signal in control^Cav2^ condition. Dashed lines mark average levels in the control^Cav2^ and grey shaded areas represent the active zone area; n in B (synapses/cultures): control^Cav2^ 50/3, cTKO^Cav2^ 48/3. **(C, D)** Quantification of the peak-to-peak distance of Munc13-1 and AP-180 (C) and of their peak levels in the periactive zone area (D). The periactive zone area is defined as an area within 68 nm on each side of the peak of the active zone marker (grey shaded areas in B); n as in B. Data are mean ± SEM; *p < 0.05, shown compared to the control^Cav2^ determined by a two-sided Student’s t-test (D for AP-180) or two-sided Mann-Whitney U tests (C,D for Munc13-1).

**Figure 4 - figure supplement 3.**
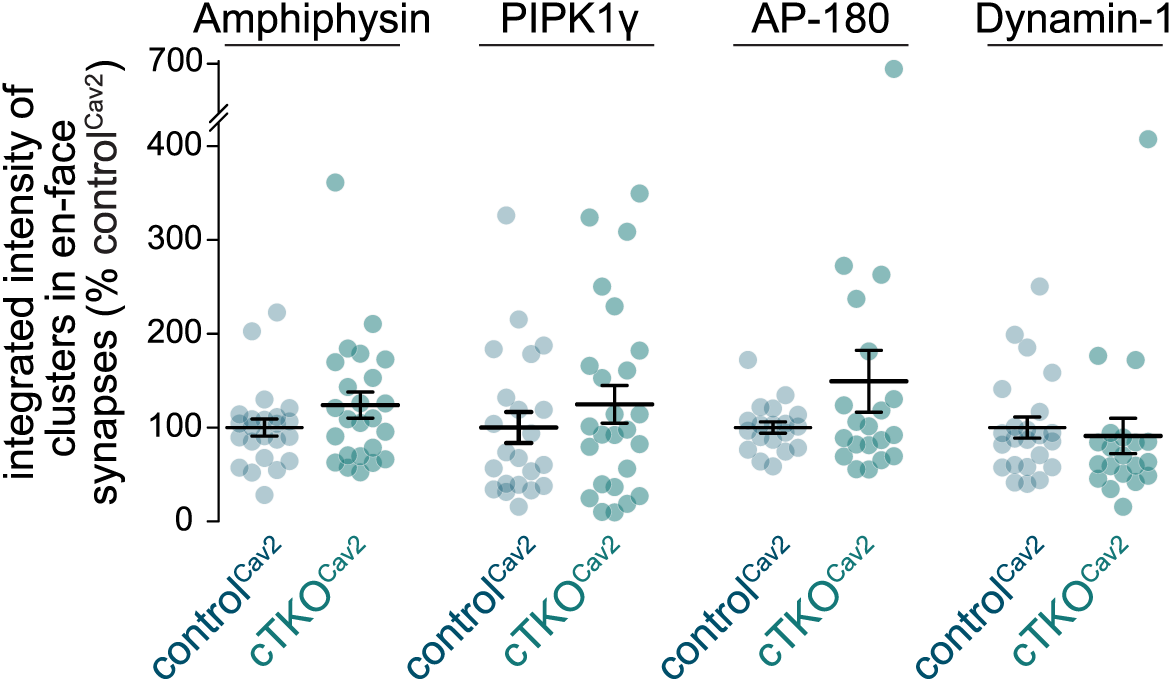
Additional analyses of en-face synapses after Ca_V_2 ablation in mouse hippocampal neurons. Quantification of the average integrated intensities (calculated as the object area multiplied by its average fluorescence intensity) of the Amphiphysin, PIPK1γ, AP-180 and Dynamin-1 objects detected in the en-face synapses quantified in Fig. 3L-Q. Intensities are normalized to the average signals in control^Cav2^ per culture; n as in Fig. 3P. Data are mean ± SEM; *p < 0.05 determined by Mann-Whitney U tests.

**Figure 5 - figure supplement 1.**
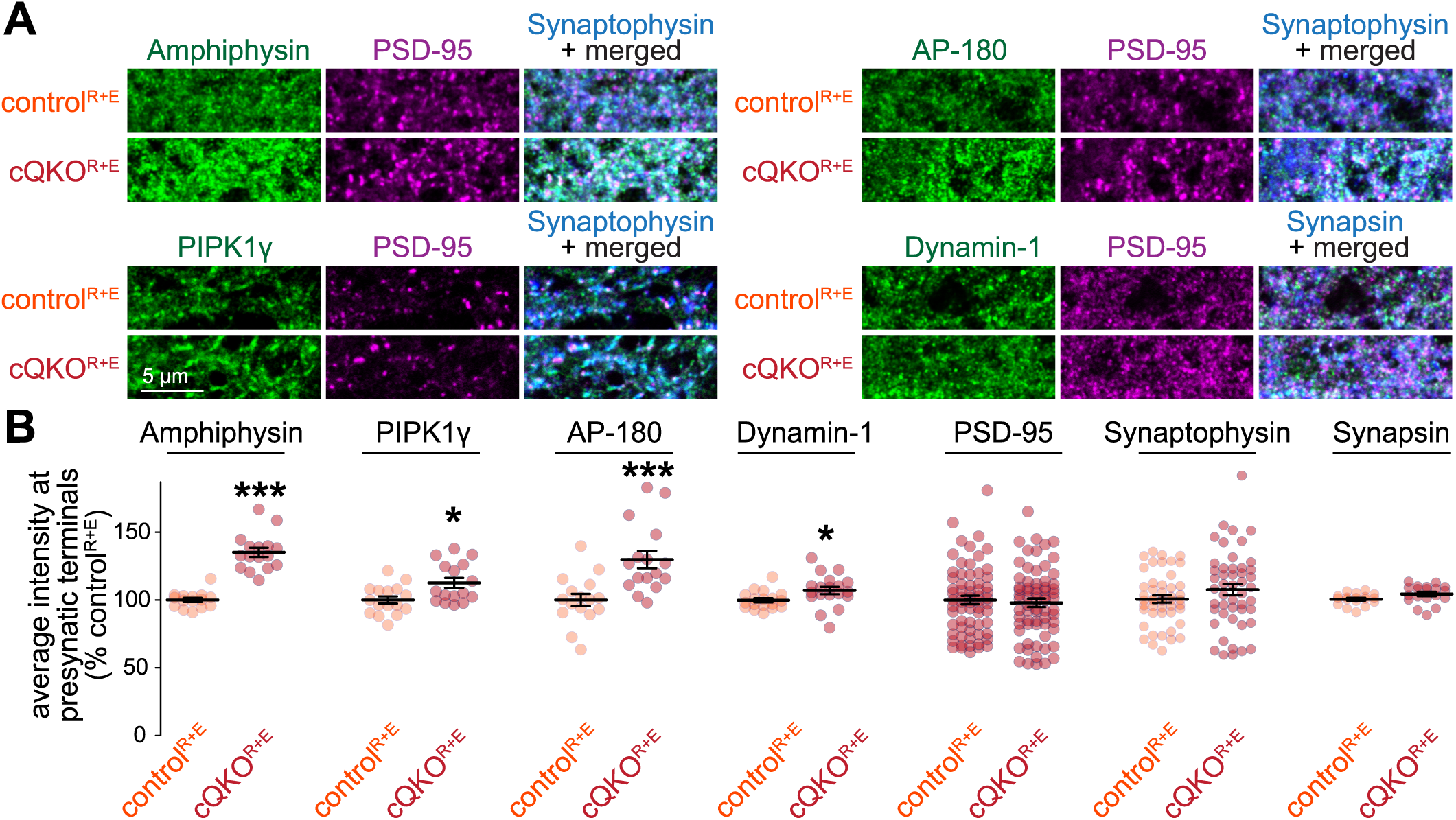
Additional analyses of endocytic proteins after active zone disruption in mouse hippocampal neurons. (**A, B**) Example confocal images (A) and quantification of the average intensities (B) of Amphiphysin, PIPK1γ, AP-180 and Dynamin-1 at synapses identified as Synaptophysin or Synapsin puncta. Intensities are normalized to the average signals in control^R+E^ per culture; n in B (images/cultures): Amphiphysin, control^R+E^ 15/3, cQKO^R+E^ 16/3; PIPK1γ, 16/3; AP-180, 16/3; Dynamin-1, control^R+E^ 18/3, cQKO^R+E^ 19/3; PSD-95, control^R+E^ 65/3, cQKO^R+E^ 67/3; Synaptophysin, control^R+E^ 47/3, cQKO^R+E^ 48/3; Synapsin, control^R+E^ 18/3, cQKO^R+E^ 19/3. Data are mean ± SEM; *p < 0.05, ***p < 0.001 determined by two-sided Student’s t tests for Dynamin-1 and Synapsin or by two-sided Mann-Whitney U tests for Amphiphysin, PIPK1γ, AP-180, PSD-95 and Synaptophysin.

**Figure 5 - figure supplement 2.**
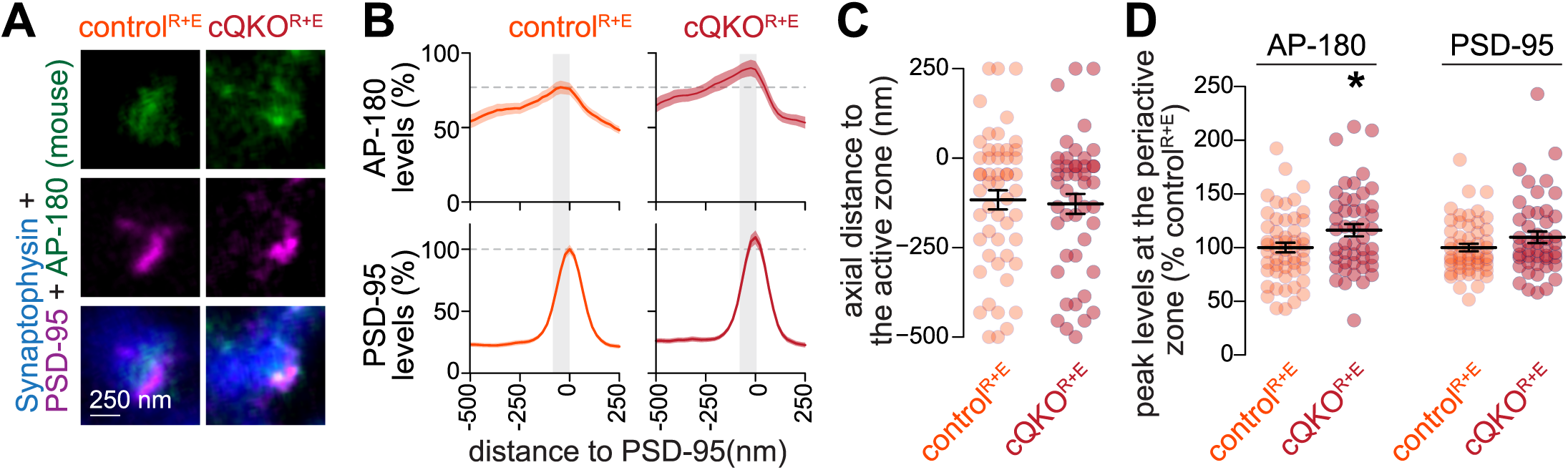
Assessment of AP-180 with an alternate antibody after active zone disruption in mouse hippocampal neurons. (**A, B**) Example side-view synapses (A) and average line profiles (B) of AP-180 (antibody A246) and PSD-95. Neurons were stained for AP-180 (imaged in STED), PSD-95 (imaged in STED), and the synaptic vesicle marker Synaptophysin (imaged in confocal). A line profile was positioned perpendicular to the center of the PSD-95 object, and synapses were aligned via the peak fluorescence of PSD-95 in the average profiles. Line profiles were normalized to the average signal in control^R+E^. Dashed lines mark average levels in the control^R+E^ condition and grey shaded areas represent the active zone area; n in b (synapses/cultures): control^R+E^ 52/3, cQKO^R+E^ 46/3. **(C, D)** Quantification of the peak-to-peak distance of PSD-95 and AP-180 (C) and of their peak levels in the periactive zone area (D). The periactive zone area is defined as the area -136 nm from the PSD-95 peak towards the presynaptic bouton (grey shaded areas in B); n as in B. Data are mean ± SEM; *p < 0.05, shown compared to the control^Cav2^ condition determined by a two-sided Student’s t-test (D for AP-180) or two-sided Mann-Whitney U tests (C, D for Munc13-1).

**Figure 5 - figure supplement 3.**
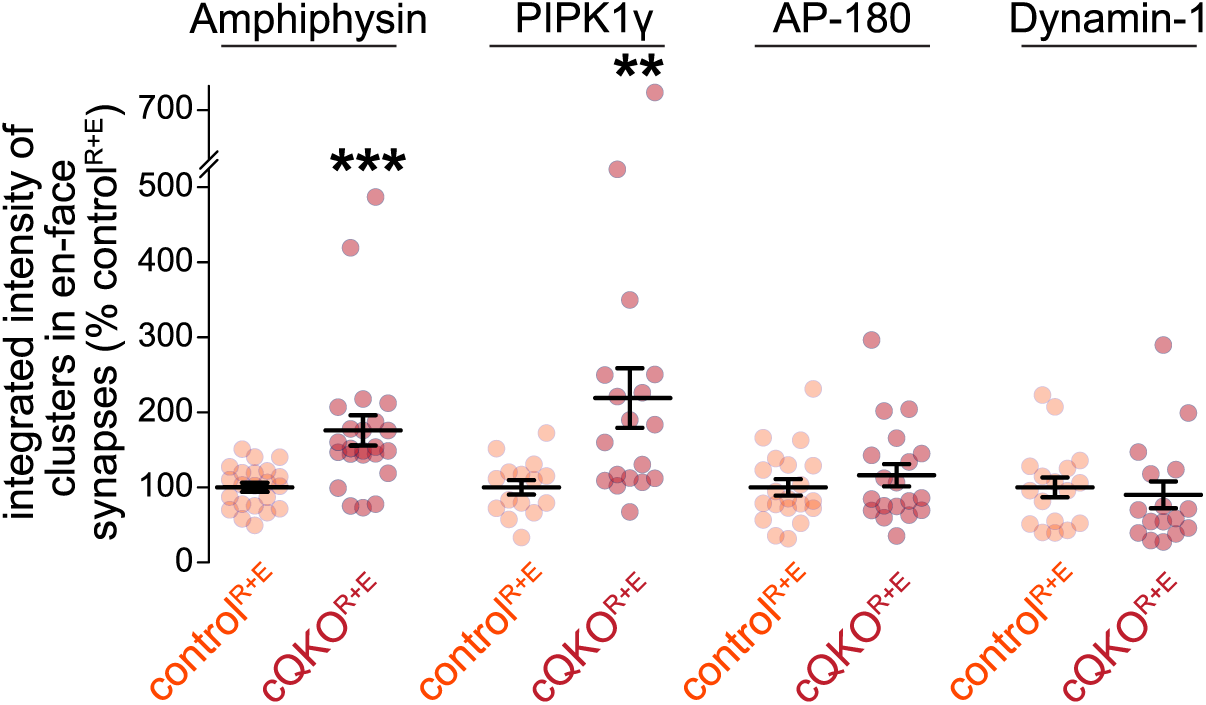
Additional analyses of en-face synapses after active zone disruption in mouse hippocampal neurons. Quantification of the average integrated intensities (calculated as the object area multiplied by its average fluorescence intensity) of the Amphiphysin, PIPK1γ, AP-180 and Dynamin-1 objects detected in the en-face synapses quantified in Fig. 5M-R. Intensities are normalized to the average signal in control^R+E^ per culture, n as in Fig. 5Q. Data are mean ± SEM; **p < 0.01 ***p < 0.001 determined by two-sided Mann-Whitney U tests.

**Figure 7 - figure supplement 1.**
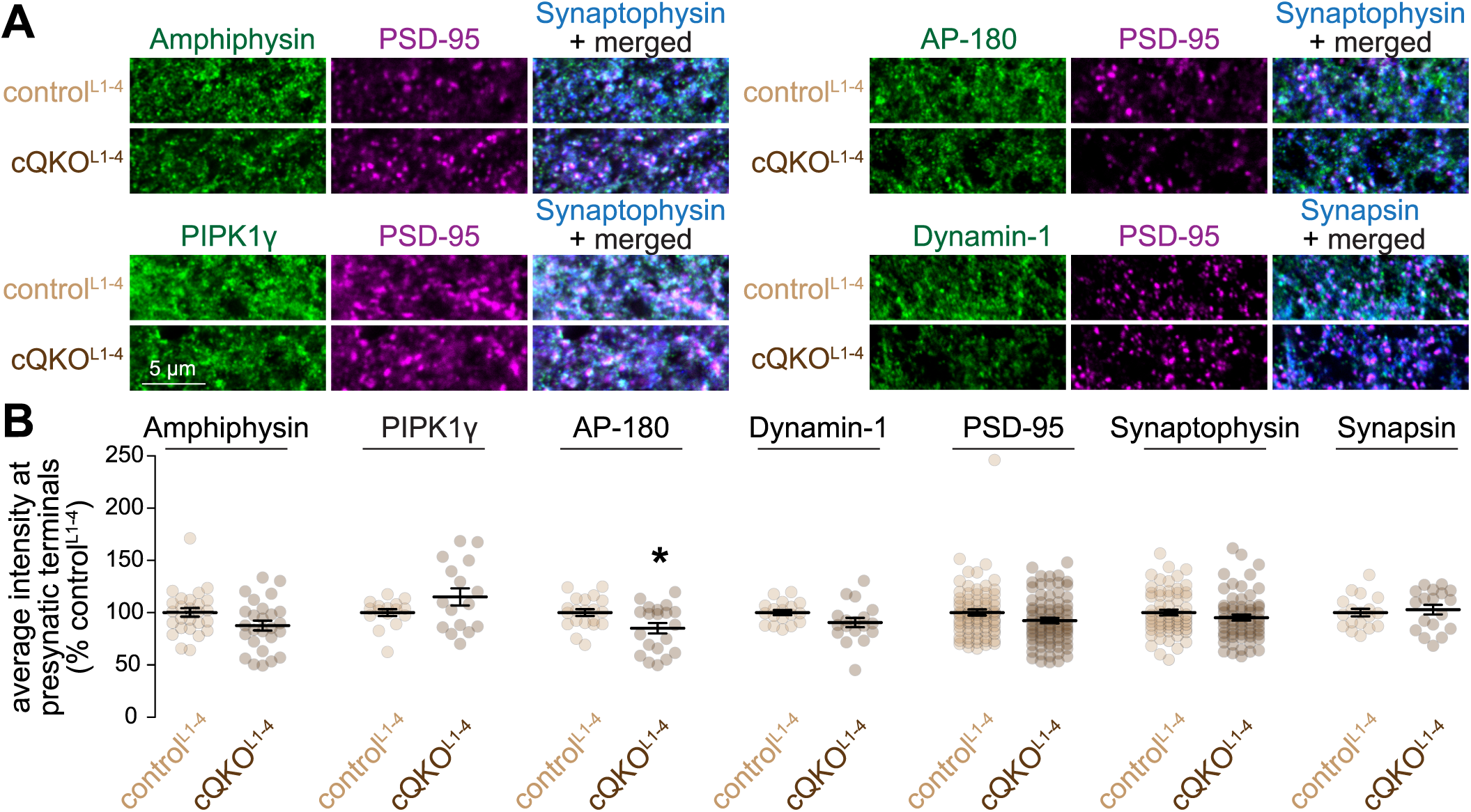
Confocal microscopic analyses of synapses after Liprin-α ablation in mouse hippocampal neurons. (**A, B**) Example confocal images (A) and quantification of the average intensities (B) of Amphiphysin, PIPK1γ, AP-180 and Dynamin-1 at synapses identified as Synaptophysin or Synapsin puncta. Intensities are normalized to the average signal in control^L1-4^ per culture; n in B (images/cultures): Amphiphysin 26/3, PIPK1γ 16/3, AP-180 20/3, Dynamin-1 18/3, PSD-95 80/3, Synaptophysin 62/3, Synapsin 18/3. Data are mean ± SEM; *p < 0.05 determined by two-sided Student’s t-tests for Synapsin and Dynamin-1 or two-sided Mann-Whitney U tests for Amphiphysin, PIPK1γ, AP-180, PSD-95 and Synaptophysin.

**Figure 7 - figure supplement 2.**
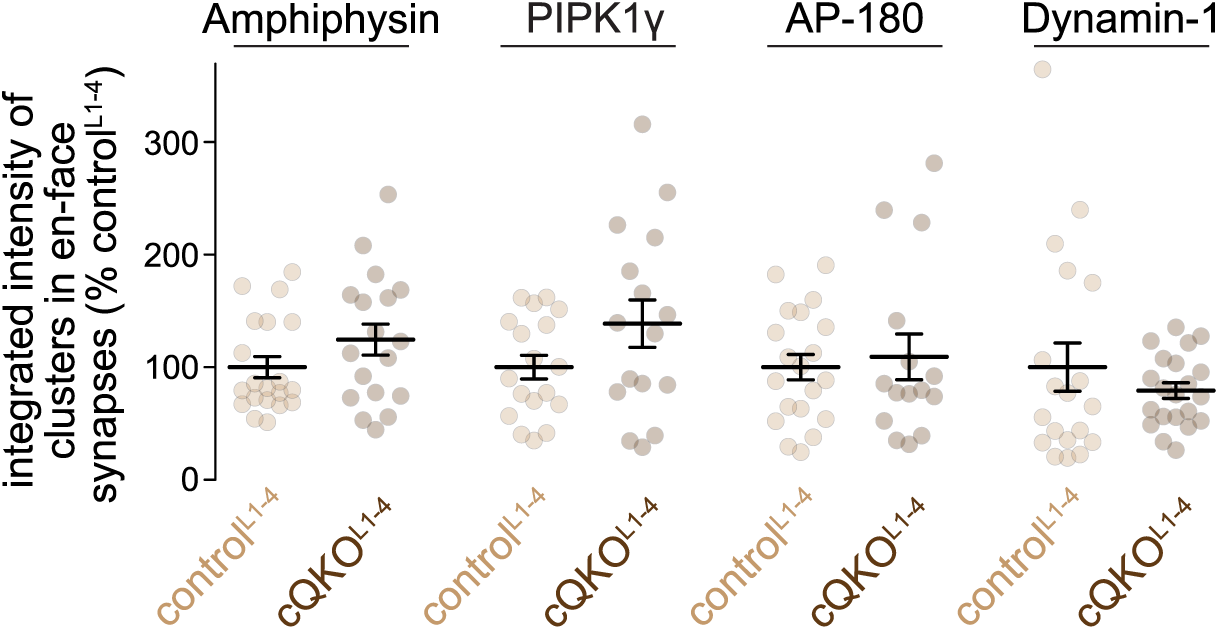
Additional analyses of en-face synapses after Liprin-α ablation in mouse hippocampal neurons. Quantification of the average integrated intensities (calculated as the object area multiplied by its average fluorescence intensity) of the Amphiphysin, PIPK1γ, AP-180 and Dynamin-1 objects detected in the en-face synapses quantified in Fig. 7M-R. Intensities are normalized to the average signals in control^L1-4^ per culture; n as in Fig. 7Q. Data are mean ± SEM; statistical significance was assessed by two-sided Mann-Whitney U tests.

